# Label free metabolic imaging to enhance the efficacy of Chimeric Antigen Receptor T cell therapy

**DOI:** 10.1101/2024.02.20.581240

**Authors:** Dan L. Pham, Daniel Cappabianca, Matthew H. Forsberg, Cole Weaver, Katherine P. Mueller, Anna Tommasi, Jolanta Vidugiriene, Anthony Lauer, Kayla Sylvester, Madison Bugel, Christian M. Capitini, Krishanu Saha, Melissa C. Skala

**Author notes:** These authors contributed equally to this work (Dan L. Pham, Daniel Cappabianca).

## Abstract

Chimeric antigen receptor (CAR) T cell therapy for solid tumors remains challenging due to the complex manufacturing process and the immunosuppressive tumor microenvironment. The manufacturing condition directly impacts CAR T cell yield, phenotype, and metabolism, which correlate with *in vivo* potency and persistence. Optical metabolic imaging (OMI) is a non-invasive, label-free method to evaluate single cell metabolism based on autofluorescent metabolic coenzymes NAD(P)H and FAD. Using OMI, we identified the dominating impacts of media composition over the selection of antibody stimulation and/or cytokines on anti-GD2 CAR T cell metabolism, activation strength and kinetics, and phenotype. We demonstrated that OMI parameters were indicative of cell cycle stage and optimal gene transfer conditions for both viral transduction and electroporation-based CRISPR/Cas9. Notably, OMI accurately predicted oxidative metabolic phenotype of virus-free CRISPR-edited anti-GD2 CAR T cells that correlated to higher *in vivo* potency against neuroblastoma. Our data supports OMI’s potential as a robust, sensitive analytical tool that enables dynamic and optimal manufacturing conditions for increased CAR T cell yield and metabolic fitness.

**One sentence summary:** Autofluorescence imaging informs manufacturing conditions that enhance yield and metabolic fitness of CAR T cells for neuroblastoma.

## INTRODUCTION

Chimeric Antigen Receptor (CAR) T cells are genetically engineered to display synthetic receptors that target antigens of interest such as tumor associated antigens and facilitate cytotoxic functions to eliminate cancer cells. FDA-approved CAR T cell treatments for hematological malignancies have demonstrated promising clinical outcomes, with patients from early trails maintaining decade-long remission (*1*). However, the translation of CAR T therapy to solid tumors remains challenging (*2*). Current CAR T manufacturing starts with (1) activation of T cells within a leukapheresis product from cancer patients using antibody stimulation and cytokine supplementation, (2) introduction of the CAR transgene, and (3) expansion of the resultant CAR T cells to reach sufficient dosage for product release and long-term treatment response (*3*). Each of these steps requires optimization for successful clinical translation of CAR T cells for solid tumors.

First, antibody activation is critical to prime T cells for proliferation and genome editing. Though the effects of certain culture media and stimulating antibodies on T cell expansion and phenotype have been characterized (*4, 5*), their synergistic impacts on CAR T cell metabolism, functions, and transgene incorporation efficiency have yet to be explored, making the selection for optimal activation conditions challenging. Second, while viral transduction is the common CAR gene transfer method used in all six FDA-approved products, it suffers from batch-to-batch variability (*6, 7*) and safety concerns due to random transgene insertion (*8*). Meanwhile, despite promising *in vivo* potency (*9, 10*), CRISPR-edited CAR T cells face challenges such as low transgene incorporation efficiency and viability (*11*). Extended *ex vivo* culture is, thus, needed to reach the desired CAR T dosage, but comes with the risk of inducing terminal differentiation that decreases *in vivo* potency (*12, 13*) while delaying patient treatments. Identifying the optimal time for CAR gene transfer could maximize yield while minimizing expansion time to increase CAR T potency and patient access. Considering remarkable metabolic changes throughout the cell cycle, T cell metabolic states following activation can potentially indicate cell cycle stage and the optimal CAR gene transfer timeframe (*11, 14, 15*). Third, media composition and cytokines used during expansion can impact CAR T cell phenotype, fitness and clinical outcomes (*16*). While several cytokine cocktails, including IL-7 and IL-15 have been studied to control CAR T cell differentiation and retain stem-like characteristics (*e.g.*, expression of CCR7, CD62L, CR45RA) for enhanced *in vivo* potency and persistence (*17, 18, 19*), metabolites in culture media such as glucose and glutamine could similarly program CAR T cell metabolic states for enhanced *in vivo* potency and persistence (*16, 18*). Metabolic fitness, such as high dependence on oxidative phosphorylation (OXPHOS) and fatty acid oxidation, low glycolytic activity, and high mitochondrial mass, has recently emerged as a key critical quality attribute for CAR T cells (*20–23*) As metabolism plays an intricate role in T cell activation, cell cycle progression, gene transfer efficiency, and in vivo potency, metabolic monitoring of T cells at various stages during CAR T manufacturing could provide complementary measurements for dynamic fine-tuning of manufacturing conditions.

Current methods for metabolic evaluation lack single cell resolution to characterize heterogeneity within the bulk T cell product (*e.g.*, metabolite quantification from media), involve manipulation of the CAR T product, and are time consuming (*e.g.*, single cell metabolomics or flow cytometry). Thus, while these methods provide critical insights for process development and optimization, their integration into a real-time, non-invasive, adaptive manufacturing workflow remains challenging. Optical metabolic imaging (OMI) is a label-free, noninvasive method to characterize metabolism within single cells (*24, 25*). OMI includes 13 metabolic parameters based on autofluorescence intensities and lifetimes of NAD(P)H and FAD, two metabolic coenzymes that are involved in hundreds of reactions in the mitochondria and cytosol (Table 1). Since reduced NAD(P)H and oxidized FAD are autofluorescent, the optical redox ratio (ORR), defined as 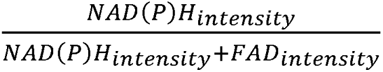can be used to measure cellular redox balance (*26–28*). Meanwhile, fluorescence lifetimes indicate the binding activity of these molecules. Due to conformational changes upon protein binding, free NAD(P)H self-quenches and has a short lifetime, while protein-bound NAD(P)H has an extended conformation and hence, longer lifetime. FAD displays an opposite trend, with free and protein-bound FAD having a long and short lifetime, respectively (*27–29*). OMI provides single cell resolution to characterize heterogeneity within population and identify important cell subsets (*24, 30*), while offering non-invasive, label-free, and real-time readouts of cell metabolism.

**Table 1.** List of 14 OMI parameters (13 metabolic features + cytoplasm size). OMI parameters were derived from autofluorescence intensity and fluorescence lifetimes of NAD(P)H and FAD. Each parameter was measured from the cytoplasmic region on a single-cell level using manually segmented masks.

| List of OMI parameters |  |
| --- | --- |
| NAD(P)H intensity | Number of NAD(P)H photons |
| NAD(P)H $\tau_1$ | Fluorescence lifetime of free NAD(P)H |
| NAD(P)H $\tau_2$ | Fluorescence lifetime of bound NAD(P)H |
| NAD(P)H $\alpha_1$ | Fraction of free NAD(P)H |
| NAD(P)H $\alpha_2$ | Fraction of bound NAD(P)H |
| NAD(P)H $\tau_m$ | Mean NAD(P)H fluorescence lifetime = $\alpha_1\tau_1 + \alpha_2\tau_2$ |
| FAD intensity | Number of FAD photons |
| FAD $\tau_1$ | Fluorescence lifetime of bound FAD |
| FAD $\tau_2$ | Fluorescence lifetime of free FAD |
| FAD $\alpha_1$ | Fraction of bound FAD |
| FAD $\alpha_2$ | Fraction of free FAD |
| FAD $\tau_m$ | Mean FAD fluorescence lifetime = $\alpha_1\tau_1 + \alpha_2\tau_2$ |
| Redox ratio | $\frac{NAD(P)H_{intensity}}{NAD(P)H_{intensity} + FAD_{intensity}}$ |
| Cytoplasm size | Number of pixels in cytoplasm mask |

Here, we test whether OMI can predict manufacturing conditions that improve CAR gene transfer efficiency and metabolic fitness of CRISPR-edited anti-GD2 CAR T cells for neuroblastoma treatment. Neuroblastoma is the most common extracranial solid tumor in children with a 5 year survival rate of less than 50% in high-risk patients (*31, 32*). Due to ubiquitous overexpression of the disialoganglioside GD2 on neuroblastoma cells, anti-GD2 CAR T cells are a promising treatment (*32, 33*); however, poor T cell persistence and potency remains. Using OMI, we characterized metabolic changes in T cells upon activation in various conditions and determined the relationship between T cell metabolism, cell cycle progression, and CAR gene transfer efficiency. We also performed OMI to determine the synergistic impacts of culture media and cytokine cocktails on CAR T cell metabolism and phenotype. Finally, OMI parameters were used to predict the expansion condition that achieved high *in vivo* potency in NOD/SCID/IL2R*γ*c^−/−^ mouse xenograft model of human neuroblastoma tumor.

## RESULTS

### OMI demonstrates the dominating impact of media composition on T cell metabolism and activation kinetics

Several medium formulations with different concentrations of key nutrients (such as glucose^high^/glutamine^high^ media ImmunoCult XF and glucose^low^/glutamine^low^ TexMACS media) are used for CAR T production in preclinical and clinical settings (*34*). Meanwhile, multiple soluble T cell activation antibodies have also been developed with different structures and compositions (such as StemCell *α*CD2/*α*CD3/*α*CD28 and TransAct *α*CD3/*α*CD28). However, the effects of these media and activating antibodies on T cell metabolism and CAR T cell generation are not fully defined. Using OMI, we characterized changes in T cell metabolism following activation by 4 unique combinations of antibody (StemCell *α*CD2/*α*CD3/*α*CD28 or TransAct *α*CD3/*α*CD28) and media (ImmunoCult XF or TexMACS) for 24, 48 and 72 hours (**Fig 1A-B**). We observed lower NAD(P)H *τ*_m_ and increased cell size following activation (**Fig 1B**). Interestingly, T cells activated in ImmunoCult XF media displayed greatest increase in the proportion of free NAD(P)H (NAD(P)H *α*_1_) at 48 hours post activation, regardless of the activating antibodies used (**Figure 1C**, right). Meanwhile, T cells activated in TexMACS media demonstrated peak increase in NAD(P)H *α*_1_ at 72 hours post activation (**Fig 1D**, right). High NAD(P)H *α*_1_ has been correlated with increased glycolysis (*35*), suggesting that T cells shift towards glycolysis upon activation, consistent with prior metabolic studies (*36*). However, our data suggest that the kinetics of metabolic changes in activated T cells depend on the media composition.

**Figure 1.**
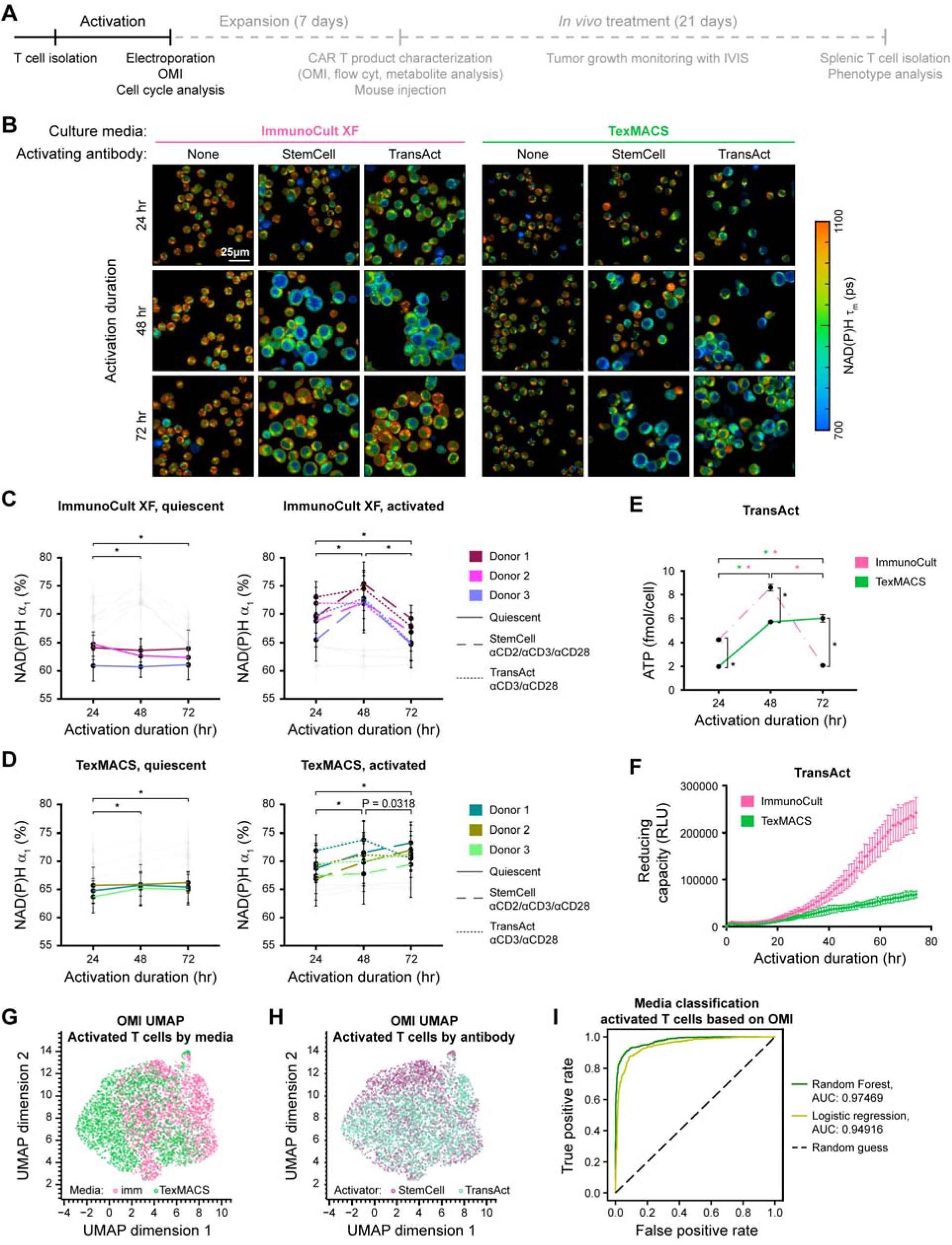
Activating media had dominating effects over activating antibody on CD3 T cell metabolism. **(A)** Experimental timeline. **(B)** Representative NAD(P)H mean lifetime (NAD(P)H) images and **(C-D)** quantification of NAD(P)H *τ*_m_ of quiescent (left) and activated (right) CD3 T cells in either **(C)** ImmunoCult XF or **(D)** TexMACS media. n = 93-290 cells/condition/donor, two-way ANOVA and adjusted multiple comparison. **(E)** ATP production and (F) reducing potential of T cells activated with TransAct *α*CD3/*α*CD28 antibody in ImmunoCult XF and TexMACS media. n = 3 replicates/condition/timepoint, two-way ANOVA with Fisher’s LSD post-hoc test for multiple comparison. **(G-H)** UMAP based on Euclidean distance projection of 14 OMI parameters (Table 1) from T cells activated for 24-72 hours, colors representing **(G)** culture media and **(H)** activating antibody, n=4923 cells from 3 independent donors. **(I)** Receiver operating characteristic (ROC) curves and areas under the curve (AUCs) Random Forest (RF) and Logistic Regression (LR) algorithms to classify T cells activated in ImmunoCult XF versus TexMACS media (regardless of activating antibody or activation duration) based on OMI parameters. Data were randomly split into 70% for training (n=3446 cells) and 30% for testing (n=1477 cells). Bars are mean ± SD. * p < 0.0001.

Metabolic reprogramming upon activation is crucial not only for energy production but also for biosynthesis to support T cell proliferation and effector functions (*37*). Thus, we next investigated the impacts of various activation conditions on ATP production kinetics and cellular reducing potentials of activated T cells. Early after activation, T cells in glucose^high^/glutamine^high^ ImmunoCult XF media exhibited significantly higher intracellular ATP levels, peaking at 48 hours, compared to those in glucose^low^/glutamine^low^ TexMACs media. However, this trend reversed at 72 hours post activation, when T cells activated in TexMACs reached peak ATP production and displayed significantly higher intracellular ATP levels than those in ImmunoCult XF (**Fig 1E**). This is consistent with NAD(P)H *α*_1_ kinetics as measured by OMI (**Fig 1D**). We also measured cellular reducing capacity, an indicator of the overall bioreactivity of intracellular NAD(P)H-dependent reductase enzymes, to further investigate NAD(P)^+^ reducing metabolic pathways (such as glycolysis, pyruvate conversion to acetyl-CoA, or the tricarboxylic cycle) (*38*). T cells activated in ImmunoCult XF media displayed a rapid increase in cellular reducing capacity, plateauing at 72 hours, while T cells activated in TexMACS media exhibited a slower and continuous rise in reducing potential up to 72 hours (**Fig 1F**). These observations underscore a distinct response of T cells activated in glucose^high^/glutamine^high^ ImmunoCult XF, characterized by a more rapid and pronounced increase in both ATP levels and reducing potential, and are consistent with OMI measurements.

Additionally, Uniform Manifold Approximation and Projection (UMAP) of OMI parameters showed clustering based on culture media rather than the type of activating antibodies used (**Fig 1G-H**), indicating that metabolic features of activated T cells were predominantly determined by media composition. Distinct OMI metabolic profiles also allowed classification of T cells activated in ImmunoCult XF media and TexMACS media with high sensitivity and specificity (AUC > 0.93) across several classifier models (**Fig 1I**). Overall, our data demonstrate that while increased glycolysis upon activation is robust across several activation conditions, media composition has critical impacts on T cell metabolic kinetics and ATP production, with faster changes induced by glucose^high^/glutamine^high^ ImmunoCult XF media, regardless of the activating antibody used.

### OMI metabolic profile indicates cell cycle progression in activated T cells

Since changes in metabolism and ATP production are directly involved in T cell proliferation, we further investigated the impacts of different activation conditions on T cell cell cycle progression. Using OMI, we characterized T cell metabolism every 12-hours following activation with StemCell *α*CD2/*α*CD3/*α*CD28 antibody in ImmunoCult XF media (Imm) or TransAct *α*CD3/*α*CD28 antibody in TexMACS media (Tex) (**Fig 2A, Fig S1**). While the signature decrease in NAD(P)H *τ*_m_ and increase NAD(P)H *α*_1_ were detected in both Imm and Tex activated T cells as early as 12 hours post activation (**Fig 2B-D**), we also observed distinct metabolic kinetics among these cells. Consistent with previous data, Imm activated T cells reached peak metabolic changes (lowest NAD(P)H *τ*_m_ and highest NAD(P)H *α*_1_) faster than Tex activated T cells (36-48 hours compared to 48-72 hours post-activation, respectively) (**Fig 2C, D**). We also observed consistently greater effect sizes (Glass’s Delta) of the Imm activation condition on NAD(P)H *τ*_m_ and NAD(P)H *α*_1_ across 3 donors throughout the 72-hour time course (**Table S1**). These findings further support that glucose^high^/glutamine^high^ ImmunoCult XF media induced not only faster but also greater metabolic changes in activated T cells compared to glucose^low^/glutamine^low^ TexMACS media. Besides metabolic changes, T cells also underwent a significant increase in cytoplasm size following activation by Imm and Tex methods, indicating that they are primed for proliferation (**Fig 2E**).

**Figure 2.**
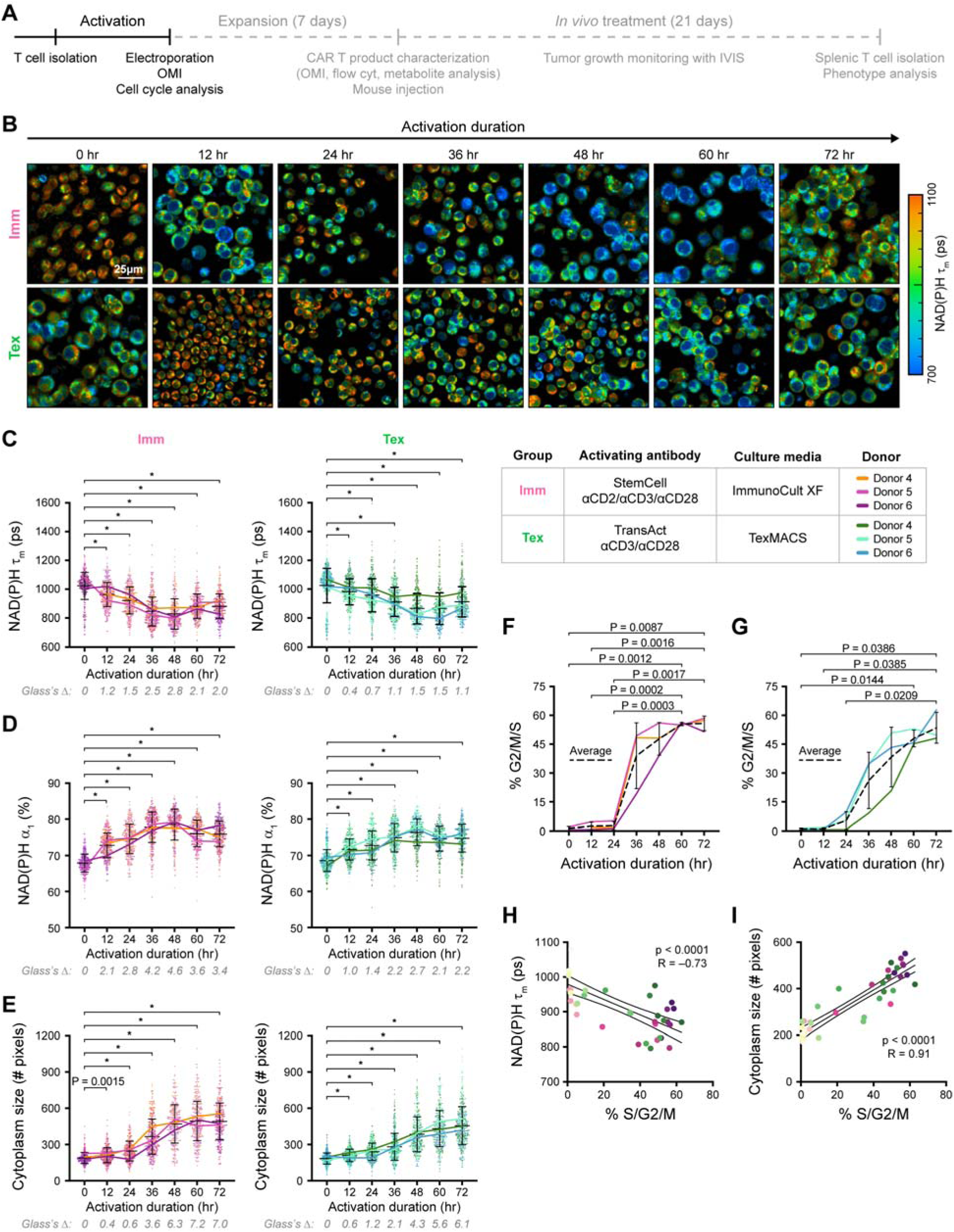
OMI is sensitive to metabolic changes as T cells progress through cell cycle following activation. **(A)** Experimental timeline. **(B)** Representative NAD(P)H *τ*_m_ images of T cells activated for 12-72 hours with Imm (top row) or Tex (bottom row) activation methods. Quantification of **(C)** NAD(P)H *τ*_m_, **(D)** NAD(P)H *α*_1_, and **(E)** cytoplasm size of T cells activated with Imm method (left column, pink) or Tex method (right column, green). n = 300-500 cells/condition from 3 donors, Brown-Forsythe and Welch ANOVA test with Dunnette T3 post hoc test. Glass’s Deltas were calculated with respect to quiescent cells as the effect sizes of Imm and Tex activation methods on OMI parameters. **(F-G)** Percentage of T cells in S/G2/M phase following activation with **(F)** Imm or **(G)** Tex method. n = 3 donors, Brown-Forsythe and Welch ANOVA test with Dunnette T3 post hoc test. **(H-I)** Correlation between OMI parameters, including **(H)** NAD(P)H *τ*_m_ and **(I)** cytoplasm size, and the percentage of cells in S/G2/M phase. n= 42 samples from 3 donors, Pearson R analysis. **(B-G)** Color solid lines represent donor averages. **(H-I)** Each dot represents one sample average, color coded based on the method (Imm or Tex) and duration of activation (0–72 hours). Bars are mean ± SD. * p < 0.0001.

Following activation with either Imm or Tex methods, T cells rapidly progressed through the cell cycle with increased DNA content measured via flow cytometry of Hoechst stain (**Fig S2B-C**). Consistent with the OMI metabolic kinetics, Imm activated T cells displayed an earlier increase in the percent of cells in S/G2/M phase (%S/G2/M) (36-48 hours, **Fig 2F, Fig S2B**) than Tex activated T cells (60-72 hours, **Fig 2G, Fig S2C**). Correlation analysis confirmed the relationship among OMI metabolic features and cell cycle progression. Across three donors and two activation methods, we found significant (p<0.0001) correlations between OMI measurements and %S/G2/M (**Fig 2H-I, Fig S2D**). %S/G2/M correlated negatively with NAD(P)H *τ*_m_ and positively with NAD(P)H *α*_1_ and cytoplasm size, suggesting that actively cycling T cells had low NAD(P)H *τ*_m_, high NAD(P)H *α*_1_, and large cytoplasm size (**Fig 2H-I, Fig S2D**). T cell proliferation (%Ki-67^+^ cells) also trended upwards throughout the 72-hour activation time course (**Fig S2E-F**). In summary, our data demonstrate that non-invasive, label-free OMI measurements are sensitive to the influence of media composition on cell cycle progression following activation, with glucose^high^/glutamine^high^ ImmunoCult XF media inducing faster metabolic changes to support earlier cell cycle entry compared to glucose^low^/glutamine^low^ TexMACS.

### OMI identifies features of T cells that predict optimal CAR gene transfer conditions

T cell activation primes them for CAR gene transfer; however, our data indicated that different activation conditions yielded different T cell metabolic profiles and cell cycle progression kinetics. Thus, we further investigated how T cell characteristics at gene transfer timeframe affected the gene transfer efficiency. Using OMI, we analyzed anti-GD2 CAR T products generated by retroviral transduction (**Fig S3**) and virus-free electroporation (EP)-based CRISPR/Cas9 **(Fig 3)**. T cells isolated from two healthy donors were activated (day 0) and underwent T-cell receptor alpha-constant knockout (*TRAC* KO) (day 2) prior to retroviral transduction with anti-GD2 CAR constructs (14G2a-OX40-CD28-*ζ* CAR or 14G2a-41BB-*ζ* CAR) (day 4) as previously described (**Fig S3A**) (*9*). At the transduction timepoint (day 4), *TRAC* KO T cells displayed significantly lower NAD(P)H *τ*_m_, higher NAD(P)H *α*_1_, and greater cytoplasm size – OMI features of cycling cells – compared to control T cells with intact T cell receptor (TCR) (**Fig S3B-E**). Importantly, we observed significantly higher transduction efficiency, quantified as percent CAR positivity post-expansion (day 11), in *TRAC* KO cells compared to control cells (**Fig S3F**). This suggests the potential of OMI metabolic features to inform transduction conditions that increases CAR yield via viral transduction.

**Figure 3.**
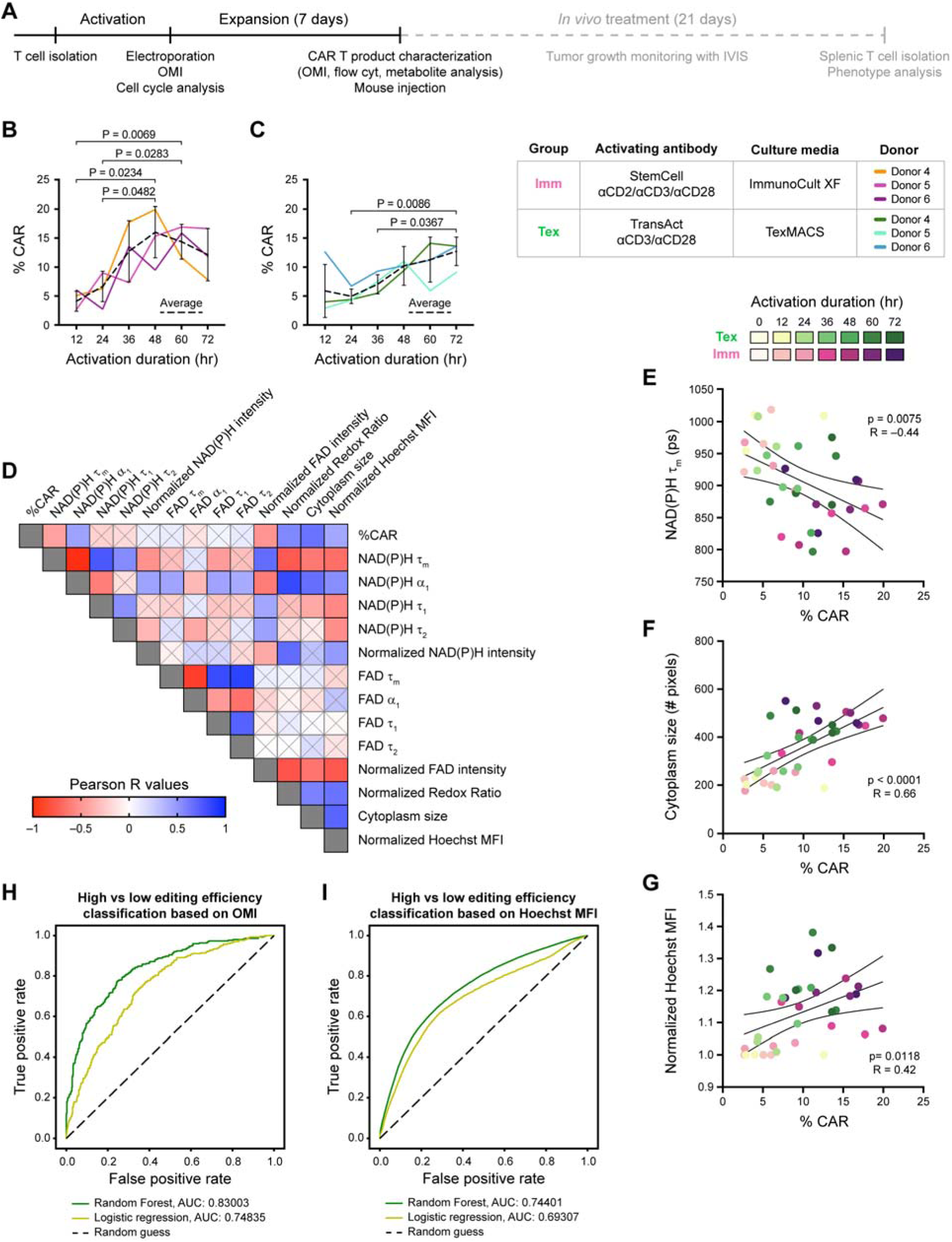
OMI identifies optimal timeframe for CRISPR/Cas9 genome editing. **(A)** Experimental timeline. **(B-C)** Genome editing efficiency (%CAR positivity) for T cells activated with **(B)** Imm and **(C)** Tex methods for 12-72 hours. n = 4-5 replicates across 3 donors, Brown-Forsythe and Welch ANOVA test with Dunnette T3 post hoc test. **(D-G)** Pearson R correlation analysis among OMI parameters, Hoechst median fluorescence intensity (MFI) at EP, and genome editing efficiency post expansion. NAD(P)H intensity, FAD intensity, redox ratio and Hoechst MFI were normalized to 12-hour-activated group averages. n = 36 samples across 3 donors. Crossed out cells represent non-significant correlations (p > 0.05). **(E-G)** Each dot represents one sample average, color coded by activation method (Imm or Tex) and duration of activation (12–72 hours). **(H-I)** ROCs and AUCs for classification models to predict gene editing outcomes [high (>15% CAR^+^ for Imm activated T cells and >10% CAR^+^ for Tex activated T cells) versus low efficiency] based on **(H)** OMI parameters and **(I)** normalized Hoechst MFIs at EP. n=3881 cells for training, 970 cells for testing for **(H)** and n = 611841 cells for training and 152960 cells for testing for **(I)**. Bars are mean ± SD. * p < 0.0001.

Besides viral vectors, CRISPR/Cas9 is currently being explored for future CAR T therapies as it allows precise genome editing that can improve treatment safety and efficacy (*39, 40*). Therefore, we next characterized whether OMI features could inform CRISPR genome editing efficiency. Following 12-to-72 hour activation by either Imm or Tex methods, T cells were imaged with OMI and electroporated to generate CRISPR-edited anti-GD2 CAR T cells as previously described (*41*). Notably, across 3 donors, the timeframe that yielded optimal genome editing efficiency, quantified as percent CAR positivity post expansion **(Fig 3B-C)**, aligned with the timeframe of maximal decrease in NAD(P)H *τ*_m_ and increase in NAD(P)H *α*_1_ **(Fig 2C-D)**. These OMI characteristics were consistent with those displayed by T cells that yielded high transduction efficiency (**Fig S3**). For Imm activated T cells, EP at 36-60 hours post-activation yielded highest CAR gene transfer efficiency (**Fig 3B**). Meanwhile, Tex activated T cells showed increased % CAR positivity as the activation duration at EP increased, with the highest genome editing efficiency observed for EP at 48-72 hours post-activation (**Fig 3C**). EP at the optimal condition yielded 3-to 4-fold-increase in genome editing efficiency.

Overall, our data suggest a robust relationship between OMI metabolic features, specifically low NAD(P)H *τ*_m_ and high NAD(P)H *α*_1_, and CAR transgene incorporation across several gene transfer platforms, including retroviral transduction and EP-based CRISPR/Cas9. This supports OMI’s potentials to inform gene transfer conditions that maximize CAR T cell yield.

### OMI features of T cells are more sensitive and specific than cell cycle analysis by flow cytometry to predict genome editing efficiency

Correlation analysis revealed significant (p < 0.05) and moderate (|R| > 0.4) correlations between genome editing efficiency (percent CAR positivity) post expansion and several OMI measurements at EP (**Fig 3D-F, Fig S3B-C**). Hoechst median fluorescence intensity (MFI) of activated T cells at EP also significantly correlated with genome editing efficiency, though the R-value was slightly lower than those of OMI variables [R = 0.42 (**Fig 3G**) compared to R = -0.44 and R = 0.66 (**Fig 3E-F**)]. Gating based on Hoechst MFI yielded the percentage of cells in S/G2/M phase, which also correlated with %CAR positivity 7 days later (**Fig S3D**). Proliferative capacity (%Ki-67^+^) in T cells also showed significant (p < 0.0001) and strong (R *≥* 0.65) positive correlations with genome editing outcome (**Fig S3E**). However, unlike label-free non-invasive OMI, S/G2/M and Ki-67 measurements by flow cytometry are destructive to samples of these cells and involve manipulating the culture during expansion.

To determine whether label-free OMI measurements could inform EP conditions, we used machine learning to classify cells from samples with high (>15% CAR^+^ for Imm activated T cells and >10% CAR^+^ for Tex activated T cells) versus low editing efficiency. A Random Forest classifier achieved high sensitivity and specificity (AUC = 0.83) in predicting genome editing efficiency based on OMI parameters at EP (**Fig 3H**). Models trained based on Hoechst MFI to predict optimal EP conditions did not perform as well (AUC = 0.74) despite 150x more cells used for training (**Fig 3I**). Overall, T cell metabolism and morphology, as quantified by non-invasive label-free OMI, are sensitive to inform the optimal EP conditions for increased CAR yield.

### Media composition determines CAR T cell OMI metabolic features and phenotypes

Given the time-intensive nature of CAR T cell expansion and its influence on T cell differentiation fate, we evaluated the synergistic impact of two media compositions (glucose^high^/glutamine^high^ ImmunoCult XF media and glucose^low^/glutamine^low^ TexMACS media) and various cytokine combinations (IL-2, IL-7, IL-7/IL-2, and IL-7/IL-15) on CRISPR-edited anti-GD2 CAR T cells. Both heatmap and UMAP of OMI measurements from CAR T cell products post-expansion (which included both CAR^+^ and CAR^-^ cells) revealed distinct clustering based on culture media rather than supplemented cytokines (**Fig 4B-D**). Notably, CAR T cells expanded in glucose^low^/glutamine^low^ TexMACS media exhibited lower FAD mean lifetime (FAD *τ*_m_), regardless of cytokines used (**Fig S5B**). Machine learning algorithms achieved high sensitivity and specificity in classifying CAR T cells expanded in glucose^high^/glutamine^high^ ImmunoCult XF media and glucose^low^/glutamine^low^ TexMACS media based on their OMI features, further confirming their distinct metabolic profiles (AUC > 0.96) (**Fig 4E**). Similar trends were observed in the CAR^+^ fraction with clustering based on culture media in a UMAP of OMI parameters (**Fig S5C-D**) and high accuracy for classification of CAR^+^ cells by media condition based on OMI features (**Fig S5E**). We did, however, observe consistently higher NAD(P)H *τ*_m_ in CAR T cells expanded in IL-7±IL-15 compared to those expanded in IL-2 in ImmunoCult XF media (**Fig S5F**), suggesting that CAR T cells did modulate their metabolism based on the cytokines present, though to a lesser extent than the culture media.

**Figure 4.**
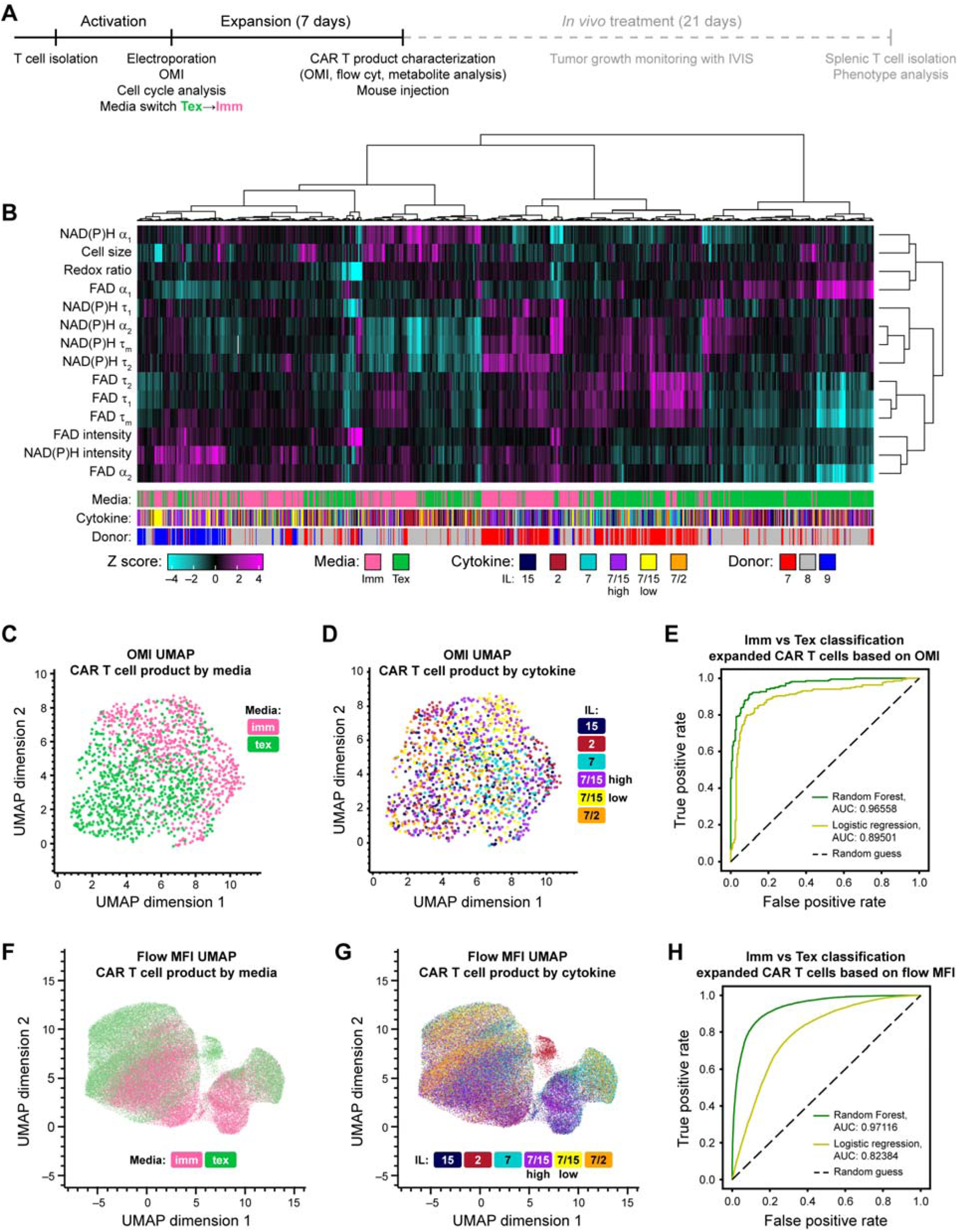
Media composition has a significant impact on CAR T cell metabolism and phenotype post-expansion. **(A)** Experimental timeline. **(B)** Z-score heatmap based on 14 OMI parameters of CAR T cells (both CAR+ and CAR-) expanded in ImmunoCult XF (Imm) or TexMACS (Tex) media with various cytokines. Hierarchical clustering was determined based on Euclidean distance from population mean using Ward’s criterion. **(C)** UMAP based on Euclidean distance of OMI metabolic parameters for CAR T cells at the end of manufacturing, color coded by **(C)** culture media or **(D)** supplemented cytokines. n = 1417 cells from 3 donors for (**B-D**). **(D)** ROC curves and AUCs of models to classify CAR T cells expanded in ImmunoCult XF or TexMACS media based on OMI parameters. n= 1134 cells for training and n = 283 cells for testing. **(E-F)** UMAP based on Euclidean distance of surface marker median fluorescence intensity (MFI) (CD27, CD45RO, CD62L, CD28, CD45RA and CCR7). n = 87317 cells across 3 donors. **(G)** ROC curves and AUCs of RF and LR classifiers to classify T cells expanded in ImmunoCult XF or TexMACS media based on surface marker expression. n = 52390 cells for training n = 34927 cells for testing.

Importantly, media composition also had dominating impacts on CAR T cell phenotype (**Fig 4F-H**). Flow cytometric analysis revealed significantly higher CCR7 expression and lower CD62L expression in CAR T cells expanded in glucose^low^/glutamine^low^ TexMACS compared to those expanded in ImmunoCult XF media (**Fig S6B-C**). UMAP based on surface marker expression also showed distinct clusters of CAR T cells expanded in ImmunoCult XF versus TexMACS media, regardless of cytokines used **(Fig 4F-G)**. Consistently, machine learning algorithms trained on surface marker expression to classify CAR T cells expanded in ImmunoCult XF versus TexMACS media also achieved high sensitivity and specificity (AUC = 0.97) (**Fig 4H**). Interestingly, CAR T cells expanded in IL-2 showed a distinct phenotypic island that was not observed in other cytokine conditions (red circle, **Fig S6E**). However, the impact of cytokines on CAR T cell phenotype was still secondary compared to culture media composition, shown by overlapping clusters of cytokine conditions in the OMI and surface marker UMAPs (**Fig 4D**, **Fig 4G**).

### OMI reveals distinct metabolic profile of CAR T cells undergoing media switch from TexMACS to ImmunoCult XF

Transient glucose or glutamine restriction has been shown to produce CAR T cells with higher metabolic fitness, characterized by greater mitochondrial mass and more oxidative activity, that correlates to better *in vivo* potency and persistence (*18, 42, 43*). As CAR T cells expanded in glucose^high^/glutamine^high^ ImmunoCult XF and glucose^low^/glutamine^low^ TexMACS exhibited different phenotypic and OMI profiles (**Fig 4**), we further investigated how changing expansion condition from TexMACS + 10ng/mL IL-7 pre-EP, which served as a transient glucose and glutamine restriction, to ImmunoCult XF + 500U/mL IL-2 post-EP (Tex➔Imm) affected CAR T cell metabolic profile and function (**Fig 5, Fig S7**).

**Figure 5.**
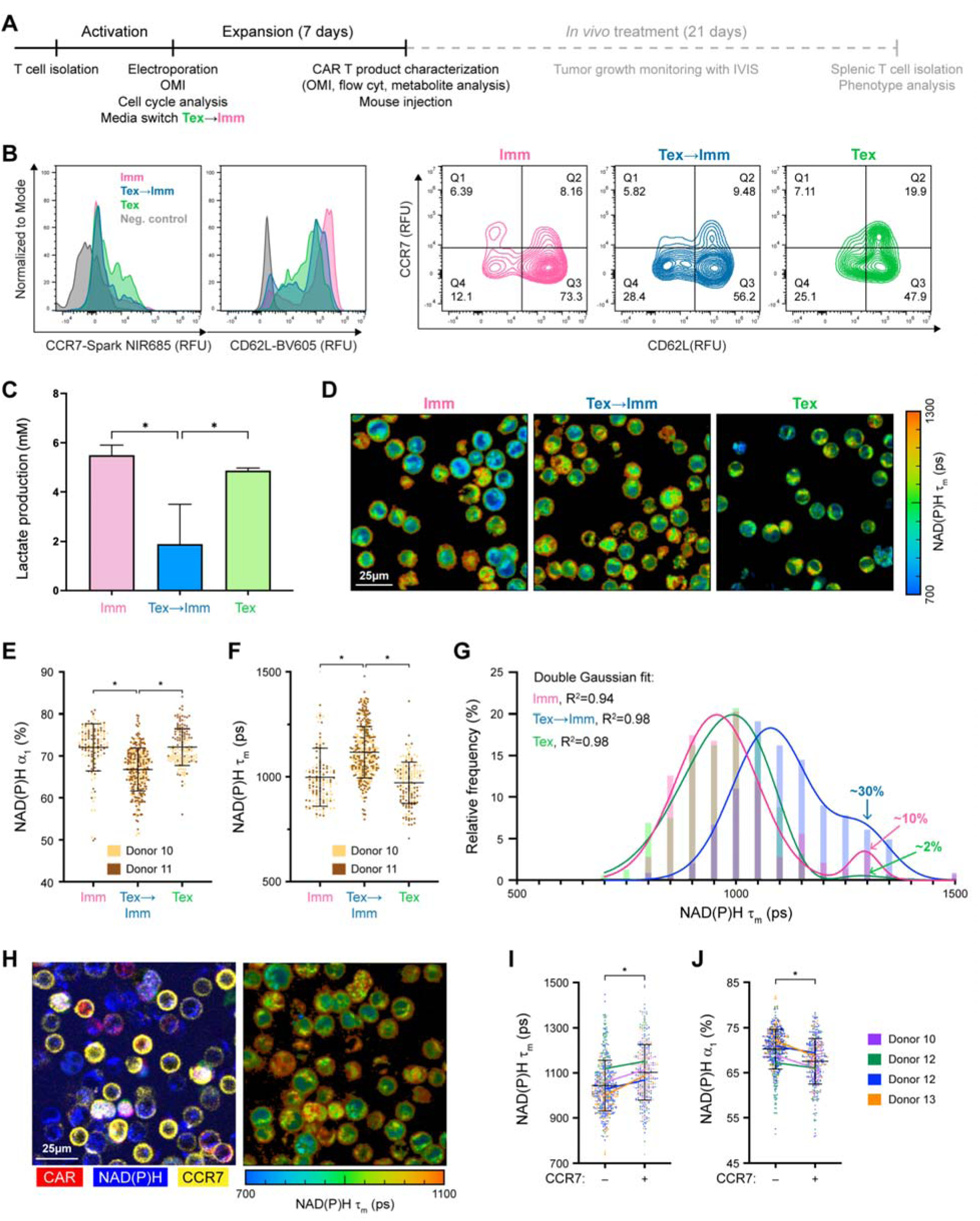
CAR T cells undergoing media transition showed a distinct OMI metabolic profile with lower glycolytic activity compared to those expanded in singular media. **(A)** Experimental timeline. **(B)** CCR7 (left) and CD62L (right) expression profile of pre-infusion CAR T cells. **(C)** Lactate production from 1 million CAR T cells within 24 hours. n = 3-15 samples/condition, Brown-Forsythe and Welch ANOVA test with Dunnette T3 test for multiple comparisons against Tex➔Imm group. **(D)** Representative NAD(P)H *τ*_m_ images and quantification of **(E)** NAD(P)H *α*_1_ and **(F)** NAD(P)H *τ*_m_ of CAR T cells expanded in Imm, Tex, or Tex➔Imm media conditions. Brown-Forsythe and Welch ANOVA test with Games-Howell’s test for multiple comparisons. **(G)** NAD(P)H *τ*_m_ histogram of CAR T cells post expansion. R^2^ value represents goodness of fit to a double Gaussian distribution. Percentages of cells in the high NAD(P)H *τ*_m_ gaussian are noted. **(E-G)** n = 143-345 cells/condition from 2 donors. **(H)** Representative immunofluorescence and NAD(P)H *τ*_m_ image and quantification of **(I)** NAD(P)H *τ*_m_ and **(J)** NAD(P)H *α*_1_ from CCR7^+^ and CCR7^-^ CAR T cells expanded in the Tex➔Imm media condition. n = 371-519 cells/group across 4 donors, Mann-Whitney test. Bars are mean ± SD. * p < 0.0001.

Post-expansion and pre-infusion into a mouse xenograft model (**Fig. 5A**), we did not observe distinct T cell phenotypes in CAR T cells expanded in the Tex➔Imm condition compared to those expanded ImmunoCult XF (Imm) and TexMACS (Tex) media alone. CAR T cells expanded in Tex➔Imm displayed an intermediate phenotype based on naïve and stem memory markers (**Fig 5B, Fig S7B**). However, we observed significantly less lactate secreted by Tex➔Imm CAR T cells, suggesting lower glycolytic activity, compared to CAR T cells cultured in a singular media composition (**Fig 5C**). This was further supported by metabolic flux analysis that showed a significantly higher oxygen consumption rate/extracellular acidification rate (OCR/ECAR) ratio in non-edited T cells expanded in the Tex➔Imm condition compared to those cultured in Tex media alone. High OCR/ECAR ratio indicated an increased dependence on oxidative metabolic pathways in Tex➔Imm group (**Fig S7C**).

OMI captured a distinct metabolic profile in CAR T cells expanded in Tex➔Imm, which had significantly lower NAD(P)H *α*_1_ and higher NAD(P)H *τ*_m_ than CAR T cells cultured in Imm or Tex media alone (**Fig 5E-F**). As OMI allows single-cell measurements, the population heterogeneity in CAR T cells expanded within the Imm, Tex, and Tex➔Imm condition was further investigated. We observed two subpopulations in each CAR T cell group post-expansion, displaying high-and low-NAD(P)H *τ*_m_ respectively (**Fig 5G**). Notably, the high NAD(P)H *τ*_m_ subpopulation made up a greater proportion in Tex➔Imm CAR T cells (∼30-40%) compared to the other two media conditions (10% in Imm and <5% in Tex CAR T cells). Additionally, independent OMI analysis of CCR7^+^ T cells from 4 donors showed significantly higher NAD(P)H *τ*_m_ and lower NAD(P)H α_1_ than CCR7^-^ cells (**Fig 5H-J**). This suggests a potential shift towards stem-like metabolism in Tex➔Imm CAR T cells, which had more cells in the high-NAD(P)H *τ*_m_ subset (**Fig 5G**, **Fig 5I**). Overall, the distinct OMI metabolic profile of Tex➔Imm CAR T cells correlates to a lower glycolytic rate and higher dependence on oxidative pathway that can potentially improve *in vivo* persistence and potency (*20, 42*).

### CAR T cells undergoing media switch from TexMACS to ImmunoCult XF yielded better *in vivo* potency

NSG mice with established GD2^+^ CHLA-20-Luciferase neuroblastoma xenografts were treated with 4 million CAR^+^ T cells expanded under Imm, Tex, or Tex➔Imm media conditions (**Fig 6A**). By day 17 post CAR T treatment, CHLA-20 bioluminescence increased significantly in cohorts treated with CAR T cells expanded in ImmunoCult XF or TexMACS media alone, indicating tumor progression (**Fig 6B-C**). Notably, CAR T cells expanded in Tex➔Imm condition led to tumor regression (fold change in tumor flux < 1) in 2 out of 4 mice, and better control of tumor flux overall (**Fig 6B-C, Fig S7D-E**). Interestingly, CD45+ human T cells isolated from the mouse spleen 21 days post treatment with Tex➔Imm CAR T cells showed better retention of stem cell memory phenotype (CCR7^+^/CD62L^+^) and resistance to exhaustion *in vivo* (PD-1^+^/TIGIT^+^) compared to CAR T cells expanded by singular media (**Fig 6D-E**) (*43, 44*).

**Figure 6.**
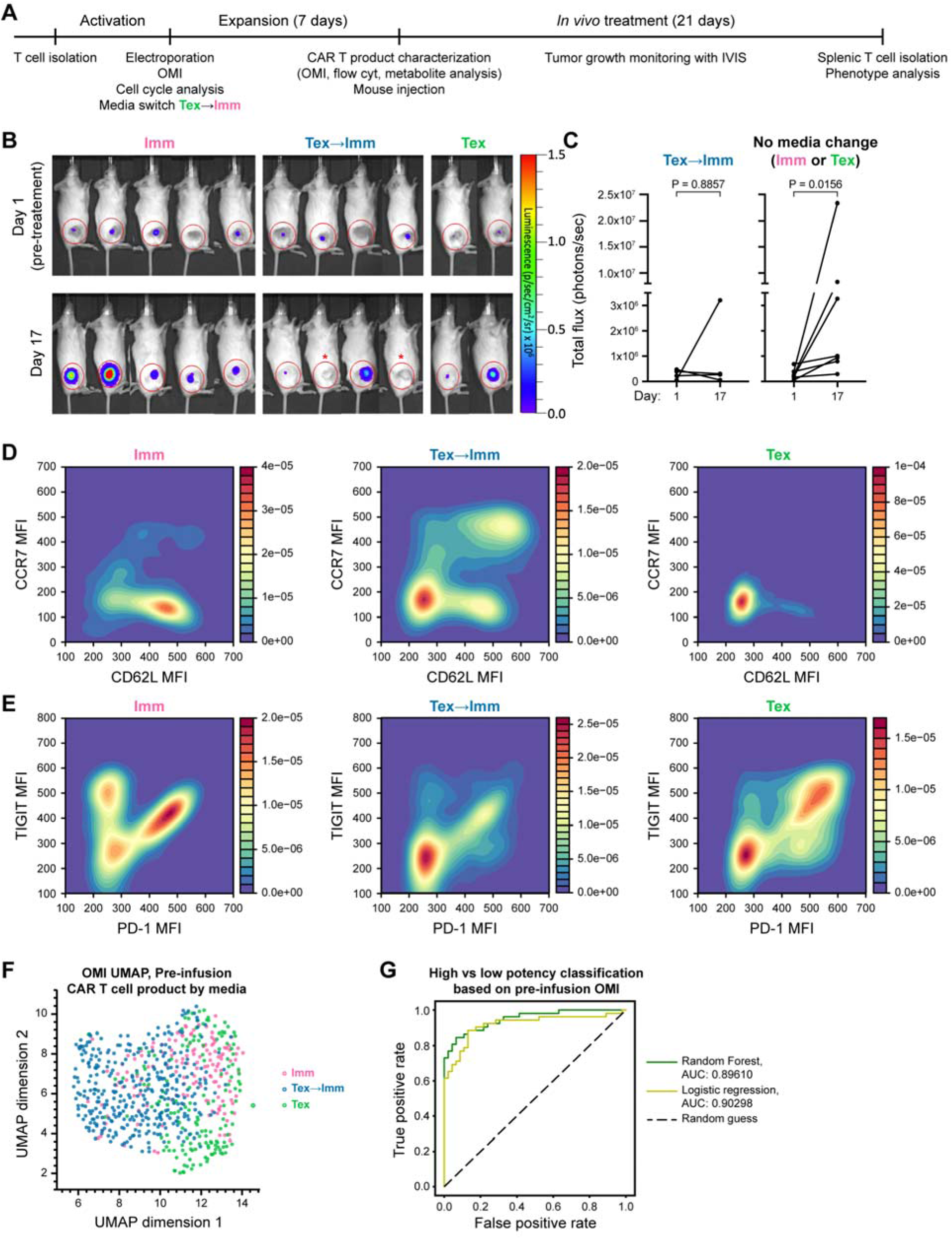
Pre-infusion OMI predicts CAR T cells with higher efficacy, preserved stem central memory phenotype (CCR7^+^ CD62L^+^) and resistant to exhaustion (PD-1^+^ TIGIT^+^) *in vivo.* **(A)** Experimental timeline **(B)** Representative images of NSG mice bearing GD2+Luciferase CHLA-20 tumors before (top row) and 17 days after (bottom row) intravenous injection with 4 million CAR^+^ T cells/mouse. Red asterisks represent tumor-free mice on day 17. **(C)** Tumor flux in NSG mice before (day -1) and after (day 17) treatment with Imm, Tex➔ Imm, and Tex CAR T cells. n= 2-5 mice/condition across 2 donors, paired non-parametric t-test (Wilcoxon test). **(D-E)** Tex➔Imm CAR T cells (both CAR+ and CAR-) expressed stem central memory markers and showed lower exhaustion *in vivo*. 2D kernel density estimation plots based on MFIs of surface markers **(D)** CCR7 and CD62L and **(E)** PD-1 and TIGIT of CAR T cells isolated from mouse spleen after 21 days of treatment. n = 9569 cells from 11 mice across 2 donors. **(F)** UMAP projection based on Euclidean distance metric of OMI parameters for CAR T cells that were activated and expanded in Imm, Tex, or Tex➔Imm conditions. n = 648 cells from 2 donors. **(G)** ROC curves and corresponding AUCs of 2 models to classify high potency (Tex➔Imm) versus low potency (Imm and Tex groups) CAR T cells based on OMI parameters of pre-infusion products. n = 518 cells for training and n = 130 cells for testing. * p < 0.0001.

We hypothesized the distinct metabolic profiles expressed by Tex➔Imm CAR T cells pre-infusion underlie their persistence and efficacy *in vivo*. To test this, we investigated whether OMI could predict CAR T *in vivo* potency based on pre-infusion metabolic features. A UMAP of OMI parameters (**Table 1**) from pre-infusion CAR T products revealed different clusters of Tex➔Imm expanded CAR T cells versus CAR T cells expanded in singular media (**Fig 6F**).

Machine learning classification based on pre-infusion OMI metabolic profiles successfully distinguished high potency (Tex→Imm) from low potency (Imm or Tex alone) CAR T cells with high sensitivity and specificity (**Fig 6G**; AUC = 0.90). These findings align with previous research indicating better *in vivo* potency by stem cell memory T cells and suggest that OMI measurements of CAR T cells *ex vivo* may predict *in vivo* response (**Fig 6G**).

## DISCUSSION

The translation of CAR T cell therapies to solid tumors is challenging, partially due to a complex manufacturing process that requires optimization at several stages. Current methods to assess T cell function are invasive (involve handling/sampling an active cell culture) and mainly rely on final product parameters to satisfy release criteria, thus, do not capture the dynamic and heterogenous changes in CAR T cells throughout manufacturing. This limits the ability to fine-tune manufacturing conditions to achieve the optimal CAR T cell profile. Recent research suggests that T cell metabolism correlates with function, and metabolic reprogramming *ex vivo* can achieve desirable functions *in vivo* (*20, 22, 42, 43*). Here, we have demonstrated that OMI, a label-free optical imaging technique, can non-invasively monitor T cell metabolism during CAR T cell manufacturing to predict conditions that improves CAR T cell products.

ImmunoCult XF and TexMACS are two common T cell culture media; however, they have different concentrations of key nutrients that impact T cell metabolism and functional fates. Specifically, ImmunoCult XF media has higher glucose and glutamine concentrations, while TexMACS media contains GlutaMax, which can be hydrolyzed into L-glutamine and L-alanine by cell-secreted aminopeptidases (*46*). Interestingly, activation in glucose^high^/glutamine^high^ ImmunoCult media induced faster and greater changes in T cell metabolism, ATP production and cellular reducing potential. Meanwhile, CAR T cells expanded in glucose^high^/glutamine^high^ ImmunoCult XF and glucose^low^/glutamine^low^ TexMACs media also displayed distinct metabolic and phenotypic profiles. Our data imply that nutrient availability and media composition modulates the strength and speed of activation, while also controlling cell cycle entry and differentiation fate. Hence, it is important to optimize CAR T manufacturing timing and media formulation to achieve the desirable product profile. Furthermore, our data reveal a robust relationship between T cell metabolism, measured by OMI, and cell cycle stage, as well as gene transfer efficiency across various platforms (viral transduction and EP-based CRISPR/Cas-9). These findings indicate that OMI could identify optimal gene transfer conditions to increase CAR yield.

CAR T cells undergoing media switch from TexMACS pre-EP to ImmunoCult XF post-EP exhibited OMI profiles indicative of high OXPHOS and low glycolytic activity while displaying improved *in vivo* potency. This suggests that modulating T cell metabolism based on different energy requirements at several manufacturing stages can improve CAR T efficacy, and is consistent with previous studies on the relationship between glycolysis and OXPHOS metabolism and T cell exhaustion and persistence (*47–49*). Metabolic reprogramming of CAR T cells for solid tumors is, therefore, of critical important, to prime them for the metabolically-stressed, immunosuppressive tumor microenvironment during tumorigenesis and metastasis (*48, 49*). We also identify a substantial sub-population with higher NAD(P)H *τ*_m_ in CAR T cells expanded in Tex➔Imm that potentially contributes to their increased *in vivo* potency and persistence. These observations emphasize the significance of single-cell resolution in characterizing population heterogeneity to identify therapeutically beneficial subsets.

There are important caveats to this study. T cells isolated from healthy donors were used as starting material for CAR T cell manufacturing, which does not fully capture characteristics observed in autologous T cells from cancer patients. Furthermore, the absence of endogenous TCR in our virus-free CRISPR TRAC-KO anti-GD2 CAR T model may influence their response to media and cytokines. Further research should explore the effects of media composition and cytokines on CAR T cells with an intact TCR and validate the relationship between OMI measurements and *in vivo* potency across various CAR T cell and tumor models. Additionally, the current two-photon laser scanning OMI system can be adapted into other configurations such as a single-photon microscope, microfluidic devices (*51*), or flow cytometer (*52, 53*) to achieve faster speed, automation, and a smaller footprint to better integrate into CAR T manufacturing workflows. These technical efforts will expand the throughput and scope of OMI as a non-invasive, sensitive tool to support the translation of CAR T cell therapy for solid tumors.

In summary, we have demonstrated the sensitivity and specificity of OMI for measuring T cell metabolism at several stages throughout CAR T manufacturing to inform optimal genome editing timeframes and expansion conditions that enhance CAR T potency and persistence. We have established robust correlations between OMI and gene transfer efficiency by retroviral vectors or EP-based CRISPR and validated OMI measurements with standard metabolic assays such as metabolite plate-based assays and extracellular flux analysis. These findings support the applications of OMI to address the current technology needs in CAR T manufacturing (*54*).

## MATERIALS AND METHODS

### Study Design

Sample size for each *in vitro* experiment includes at least 3 donors with at least 50 cells/ experimental group/donor to capture intra- and inter-donor heterogeneity based on a previous power analysis for the sensitivity of OMI to T cell activation (*35*). For the mouse studies, two donors were used across two independent experiments and the number of mice per group was determined by prior study on the same virus-free CRISPR-edited anti-GD2 CAR T model (*9*), with at least 4 mice in each group receiving either media-transitioned (Tex➔Imm) CAR T cells or non-media-transitioned (Imm or Tex) CAR T cells. Rules for stopping data collection was based on the 10 day CAR T cell manufacturing process set by prior protocols (*9*). Mouse tumors were followed until tumor volume reached 4000mm^3^. We did not exclude any data besides the mouse studies, in which one mouse was excluded due to out-of-range starting tumor size as determined by IVIS reading. We have not otherwise removed outliers. Primary and secondary endpoints were prospectively selected, and appropriate statistical corrections were applied to multiple endpoints. The number of repeats for each experiment is given in the figure caption to show that results were substantiated across multiple donors. Replication is performed at several levels: cell, image, and donor, with 4–7 technical replicates (images) per experiment. Each main finding of the paper (OMI endpoints for classification and correlation analysis to determine impacts of media composition on T cell activation, and to predict cell cycle entry, optimal gene transfer conditions, and CAR T metabolic fitness) is supported by 650-4800 T cells from 2-3 donors. Supportive experiments (ATP production, cellular reducing potential, extracellular flux analysis, lactate measurement) include at least 3 technical replicates from 1-2 donors.

The objectives of our research were to demonstrate that OMI can determine (1) cell cycle entry timeline, (2) optimal CAR gene transfer conditions, and (3) media compositions that enhance metabolic fitness of CAR T cells. These were pre-specified objectives. Research subjects were healthy volunteers and NSG mice bearing GD2+ Luciferase CHLA-20 tumors. This was a controlled laboratory experiment where treatments and measurement techniques are described below. Mice were randomly assigned among treatment groups as described below. The mouse studies were performed by M.H.F. and the investigator performing *in vitro* analyses of CAR T infusion products (D.L.P.) was blinded to the results of these mouse studies.

### T cell activation with different activating antibody and media

CD3 T cells were isolated from peripheral blood of 3 healthy donors as previously described and plated at 1 million cells/mL. T cells were divided into 4 groups and activated with 4 unique combinations of media (ImmunoCult XF media or TexMACS media) and stimulating antibodies (25*μ*L/mL StemCell *α*CD2/*α*CD3/*α*CD28 or 10*μ*L/mL TransAct *α*CD3/*α*CD28). To focus on the specific impacts of media and activating antibodies on T cell metabolism and metabolic shift kinetics, no cytokine was supplemented to activated T cells. Activated T cells were imaged using OMI at 24, 48, and 72 hours.

### Virus-free anti-GD2 CAR T cell generation and activation time course setup

Peripheral blood was drawn from healthy donors under a protocol approved by the Institutional Review Board at the University of Wisconsin – Madison (2018-0103) and informed consent was obtained from all donors. CD3 T cells were isolated by negative selection following manufacturer’s protocol (RosetteSep Human T cell enrichment cocktail, STEMCELL Technologies). Following isolation, T cells were plated at 1 million cells/mL in either ImmunoCult XF T cell Expansion Medium (STEMCELL Technologies) supplemented with 200U/mL IL-2 (Peprotech) or TexMACS Cell Culture Medium (Miltenyi Biotec) supplemented with 10ng IL-7 (Biotechne), and stimulated with 25*μ*L/mL ImmunoCult Human *α*CD2/*α*CD3/*α*CD28 T cell Activator (STEMCELL Technologies) or 10*μ*L/mL T-cell TransAct *α*CD3/*α*CD28 (Miltenyi Biotec), respectively. These two combinations of activating antibodies and culture media yielded two activation methods, Imm and Tex respectively. T cells were activated for different durations, ranging from 12 hours up to 72 hours. The metabolic profile of activated T cells was characterized with OMI while cell cycle stage and proliferative capacity were analyzed via flow cytometry of Hoechst and Ki67 staining, respectively. Cell cycle stages were gated based on median fluorescence intensity (MFI) of Hoechst stain that reflected cellular DNA content. These activated T cells were then electroporated with CRISPR/Cas9 machinery to express anti-GD2 CAR transgene as previously described (*9*). Following EP, Imm and Tex activated anti-GD2 CAR T cells were expanded for 7 days in ImmunoCult XF T cell expansion media supplemented with 500U/mL IL-2 or TexMACS media supplemented with 10ng IL-7, respectively. After expansion, CAR T cells were harvested; percent CAR positivity was quantified via flow cytometry of 1A7 anti-14G2A antibody (National Cancer Institute, Biological Resources Branch) conjugated to APC using a Lightning Link APC Antibody Labeling kit (Novus Biologicals) to evaluate genome editing efficiency as previously described (**Fig S1**) (*9, 41*).

### Virus-free anti-GD2 CAR T cell expansion and flow cytometry characterization

T cells from 3 donors were activated with either Imm or Tex method prior to EP with CRISPR/Cas9 machinery to express anti-GD2 CAR as previously described (*9, 41*). Following EP, CAR T cells were expanded in either ImmunoCult XF or TexMACS media for 7 days in a 37C, 5% CO_2_ humidified incubator. During expansion, T cell media were supplemented with 500U/mL IL-2 (Peprotech), 10ng/mL IL-7 (BioTechne), 10ng/mL IL-7 (BioTechne)+ 10ng/mL IL-15 (BioTechne) (IL-7/IL-15 low), 10ng/mL IL-7 (BioTechne) + 1ng/mL IL-15 (BioTechne) (IL-7/IL-15 low), or 10ng/mL IL-7 (BioTechne) + 100U/mL IL-2 (Peprotech) (IL-7/IL-2). Cells were counted every 2 days and adjusted to 1 million cells/mL. CAR T cell phenotypes and metabolism were analyzed with flow cytometry and OMI after 7 days of expansion. Flow cytometry was performed on an Aurora spectral cytometer (Cytek) as previously described (*9, 41*). Briefly, T cells were stained and analyzed for expression of 16 markers (anti-GD2 CAR, TCR*αβ*, CD4, CD8, CD45RA, CD45RO, CD62L, CCR7, Human CD45, PD-1, LAG3, TIM3, CD39, TIGIT, CD27, CD5) with specific clones and concentrations as previously described (*41*). An expression histogram of each surface marker was generated based on single-cell MFIs normalized by donor-matched fluorescence-minus-one (FMO) control. For *in vivo* treatment, 3 groups of CAR T cells with different expansion conditions were generated. Imm CAR T cells were activated in ImmunoCult XF media + 200U/mL IL-2 and expanded in ImmunoCult XF media + 500U/mL IL-2 following EP. Tex CAR T cells were activated and expanded in TexMACS media + 10ng/mL IL-7. Meanwhile, Tex➔Imm CAR T cells were activated in TexMACS media + 10ng/mL IL-7 and expanded in ImmunoCult XF media + 500U/mL IL-2 after EP.

### OMI of T cells

T cells were plated on a 35mm Poly-D-Lysine glass bottom imaging dish (MatTek) at the density of 200,000 cells/75*μ*L and allowed to settle for at least 15 minutes prior to imaging. Throughout the process of OMI, T cells were kept in a stage top incubator (37C, 5% CO2) to maintain their physiological conditions. OMI was performed on a custom-built multiphoton microscope (Ultima, Bruker) consisting of an inverted microscope body (Ti-E, Nikon) coupled to an ultrafast tunable laser source (Insight DS+, Spectra Physics). Images were acquired using time-correlated single-photon counting electronics (SPC 150, Becker & Hickl GmbH) using Prairie View Software (Bruker). NAD(P)H and FAD were excited at 750 nm (2.5 mW) and 890nm (4.5mW), respectively, using a 40X water immersion 1.15 NA objective (Nikon) with 2.5x optical zoom, 4.8 µs pixel dwell time, 60s integration time, and image size of 256 x 256 pixels. NAD(P)H and FAD emission were separated from excitation light using a 720 long pass filter and collected using GaAsP photomultiplier tubes (H7422, Hamamatsu) through a 440/80 nm and 550/100nm bandpass filters, respectively. PerCP conjugated CAR antibodies were excited at 980nm, while Alexa 647 CCR7 antibody was excited at 1200nm, respectively. Fluorescence emission of PerCP and Alexa 647 were collected with 690/50nm filter. Fluorescence intensity and lifetime images of NAD(P)H and FAD, together with immunofluorescence images of surface markers, were collected for each field of view (FOV), with 3-5 representative FOVs (∼150-250 cells, ∼120 µm × 120 µm) imaged per condition. The instrument response function was measured using second-harmonic generation signal from urea crystals excited at 890 nm, with full width at half-maximum of 260 ps.

### OMI analysis

Fluorescence lifetime components were computed for each image pixel using SPCImage (v8.0, Becker and Hickl GmbH) by first thresholding the background, then the pixel-wise decay curves were fit to a biexponential model convolved with the instrument response function, using an iterative parameter optimization to obtain the lowest sum of the squared differences between model and data (Weighted Least Squares algorithm). The two-component exponential decay model is 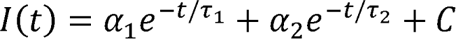, where *I(t)* is the fluorescence intensity at time *t* after the laser excitation pulse, *τ*_1_ and *τ*_2_ are the fluorescence lifetimes of the short and long lifetime components, respectively, α_1_ and α_2_ are the fractional contributions of the short and long lifetime components, respectively, and *C* accounts for background light. The mean fluorescence lifetime, *τ_m_ = α_1_τ_1_ + α_2_τ_2_,* is the weighted average of the free- and protein-bound-fraction (*55, 56*). To enhance the fluorescence counts in each decay, a bin of 1 (comprising 9 neighbor pixels) and of a bin of 2 (comprising 25 neighbor pixels) were applied on NAD(P)H and FAD lifetime images, respectively. The pixel-wise optical redox ratio was calculated as 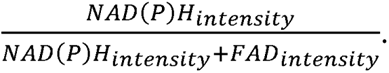 customized CellProfiler pipeline was used for manual segmentation of every individual cell nucleus within a FOV, from which the individual cell border was propagated to create a whole-cell mask. A single-cell cytoplasm mask was generated by subtracting the nuclei mask from the whole cell mask. The cytoplasm mask was applied to the corresponding OMI image to compute mean values of OMI parameters for each cell cytoplasm. 13 OMI parameters were collected and quantified (**Table 1**). Cytoplasm size was also computed for individual cells based on cytoplasm masks.

### Extracellular flux analysis

Extracellular flux assay was performed using Seahorse XF Cell Mito Stress Test Kit (Agilent). 24 hours prior to the assay, a Seahorse cell culture 96 well-plate (Agilent) was coated with 50*μ*L/well of 50 μg/mL Gibco Poly-D-lysine (ThermoFisher Scientific) for 1 hour, then washed with distilled water before storing overnight at 4C. The cell culture plate was equilibrated to room temperature before cell plating. 400,000 T cells/well were plated onto Poly-D-lysine coated cell culture plate in RPMI XF media (Agilent) supplemented with 10mM glucose (Agilent) and 2mM glutamine (Agilent) following the manufacturer’s protocol for seeding suspension cells in Seahorse XFp cell culture miniplate. Briefly, the T cell culture plate was centrifuged at 200g for 1 minute (no brake) and checked under the microscope to ensure even adhesion of T cells. T cells were then kept in a non-CO2 incubator for at least 1 hour before running the assay. 1.5*μ*M FCCP, 2.5*μ*M Oligomycin, and 0.5*μ*M Antimycin A/Rotenone were loaded into port A, B, and C respectively as metabolic inhibitors. Oxygen consumption rate (OCR) and extracellular acidification rate (ECAR) were measured with an XF96 Extracellular Flux Analyzer (Seahorse Bioscience).

### Lactate measurement

At harvest, 1 million CAR T cells for each condition were plated in 1mL fresh media (similar to their expansion condition) in a tissue culture treated 24 well-plate (VWR). After 24 hours, spent media was collected for each condition and lactate secretion was assayed in duplicate or triplicate using either Lactate Colorimetric/Fluorometric kit (abcam) or Lactate-Glo (Promega) following manufacturers’ protocols. Lactate in base TexMACS and ImmunoCult media were measured as the baseline controls. For abcam’s Lactate Colorimetric/Fluorometric kit (abcam), lactate secretion for each condition was measured based on the fluorescence intensity at 535/587nm excitation/emission using Tecan M1000 plate reader. Fitting and extrapolating of standard curve and sample measurements were performed in GraphPad Prism v10 (linear curve fitting).

### ATP production, extracellular metabolite assays, and cellular reducing capacity assay

Pan T cells were isolated (Miltenyi) from PBMCs (Leukopak, StemCell Technologies) and plated in TexMACS or ImmunoCult at 7x10^5^ cells/ml in a 6-well plate. Cells were then subsequently activated with TransAct (Miltenyi), including a non-activated control. After activation, samples were taken and incubated with RealTime-Glo™ MT (Promega) per manufacturer’s instructions in a CO_2_ and temperature-controlled plate reader (Tecan Spark Cyto). Luminescence was recorded every hour for 74 hours. Additionally, daily samples were obtained, and ATP levels were determined using CellTiter-Glo® from Promega, following the manufacturer’s instructions.

### Mouse CHLA-20 xenograft and CAR T cell treatment

All animal experiments were approved by the University of Wisconsin-Madison Animal Care and Use Committee (ACUC protocol M005915). Establishment of CHLA-20 xenograft and subsequent CAR T cell treatments were performed as previously described (*9, 41*). Briefly, male and female NOD-SCID-*γ*c^-/-^ (NSGTM) mice (9-25 weeks old; Jackson Laboratory) received 10 million AkaLUC-GFP CHLA-20 GD2^+^ human neuroblastoma cells via subcutaneous flank injection to establish tumors. Tumor flux was measured with IVIS imaging one day prior to CAR T cell treatment, and mice were divided into three treatment groups to control for equally distributed tumor flux in each group. 4 million anti-GD2 CAR^+^ cells were injected into the tail vein of tumor-bearing each mouse. Tumor flux was monitored with IVIS imaging every 3-4 days. During CAR T treatment duration, mice also received 100,000 IU of human IL-2 (National Cancer Institute, Biological Resources Branch) subcutaneously on day 0 and post imaging. Total flux was calculated using Living Image Software (PerkinElmer) as radiance (photons/second) in each pixel integrated over tumor area (cm^2^) x 4*π*. To normalize for background signal, the minimum flux value was subtracted from each image.

### Statistical analysis

All statistical analysis was performed in Prism GraphPad v10, with appropriate statistical tests chosen based on data features. p < 0.05 was chosen as the threshold for statistical significance. Multiple comparisons were adjusted with post-hoc tests. Glass’s delta for effect size was calculated as 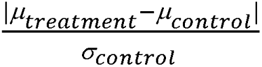 where μ_treatment_ and μ_control_ representing the means of treated and control groups, respectively; and *σ*_control_ being the standard deviation of the control group. Intensity measurements (such as NAD(P)H and FAD intensity, median fluorescence intensity (MFI) of surface markers) were normalized to donor-match control groups to account for any variations in laser power. Details on the number of sampled units and specific statistical tests for each experiment were reported in corresponding figure legends.

## List of Supplementary Materials

**Fig. S1. Activation time-course and conditions for anti-GD2 CAR T cell generation with CRISPR/Cas9.**

**Fig. S2. T cells progressed through cell cycle and proliferated upon activation.**

**Fig. S3. T cell metabolism at viral transduction timepoint correlated with transduction efficiency.**

**Fig. S4. Cell characteristics at EP correlated with genome editing efficiency.**

**Fig. S5. OMI revealed metabolic differences in CAR T cells expanded in Imm and Tex media.**

**Fig. S6. CAR+ T cells expanded in Imm and Tex media showed different phenotypes.**

**Fig. S7. CAR T cell phenotypes and metabolic profile before *in vivo* treatment course and *in vivo* treatment response.**

**Table S1. Glass’s deltas (*Δ*) showing maximal effect sizes of Imm and Tex activation methods on T cell metabolism (NAD(P)H *τ*_m_ and NAD(P)H *α*_1_).**

## Acknowledgments

We thank members of the Capitini, Saha and Skala labs for helpful discussion and comments on the manuscript, the University of Wisconsin (UW) Carbone Cancer Center Small Animal Imaging and Radiotherapy facility and Flow Cytometry Laboratory (supported by NIH P30 CA014520 and NIH S10 OD025225), Malcolm Brenner (Baylor College of Medicine) for the retroviral 14G2a-OX40-CD28-*ζ* CAR sequence and Crystal Mackall (Stanford University) for the retroviral 14G2a-41BB-*ζ* CAR sequence, the National Cancer Institute for 1A7 antibody for detection of CAR expression, Mario Otto (UW-Madison) for the parental CHLA20 cell line and James Thomson and Jue Zhang (Morgridge Institute for Research) for the AkaLUC-GFP CHLA20 cell line used for *in vivo* studies.

## Fundings

NIH R01 CA278051 (MCS, KS, CMC)

NSF Engineering Research Center (ERC) for Cell Manufacturing Technologies (CMaT) NSF-EEC 1648035 (CMC, KS, MCS)

MACC Fund (CMC)

St. Baldrick’s Foundation Empowering Pediatric Immunotherapy for Childhood Cancers Team grant (CMC, KS)

UW-Madison Office of the Vice Chancellor for Research and Graduate Education with funding from the Wisconsin Alumni Research Foundation (CMC, KS)

Hyundai Hope on Wheels (CMC, KS)

Grainger Institute for Engineering at UW-Madison (CMC, KS)

NIH R35 GM119644-01 (KS).

## Authors contributions

Conceptualization: DLP, DC, MHF, KPM, CC, KS*, MCS*

Methodology: DLP, DC, MHF, KPM, CW, KPM, AT, JV, AL, KS, MB

Data analysis: DLP, DC, MHF, CW, JV, AL, KS

Funding acquisition: CC, KS*, MCS*

Supervision: CC, KS*, MCS*

Writing – original draft: DLP, MCS*

Writing – review & editing: DLP, DC, MHF, KPM, JV, KS, CC, KS*, MCS*

* corresponding authors

## Competing interests

KS receives honoraria for advisory board membership for Andson Biotech and Notch Therapeutics. CMC receives honoraria for advisory board membership for Bayer, Elephas Bioscience, Nektar Therapeutics, Novartis, and WiCell Research Institute. MCS, DP, KS, DC declare two patents pending based on this work. No other conflicts of interest are reported.

## Data and materials availability

All data and codes used in the analysis will be available on Dryad and Zenodo.

