## Supplementary_data for "Label free metabolic imaging to enhance the efficacy of Chimeric Antigen Receptor T cell therapy"

**Supplementary Figures**

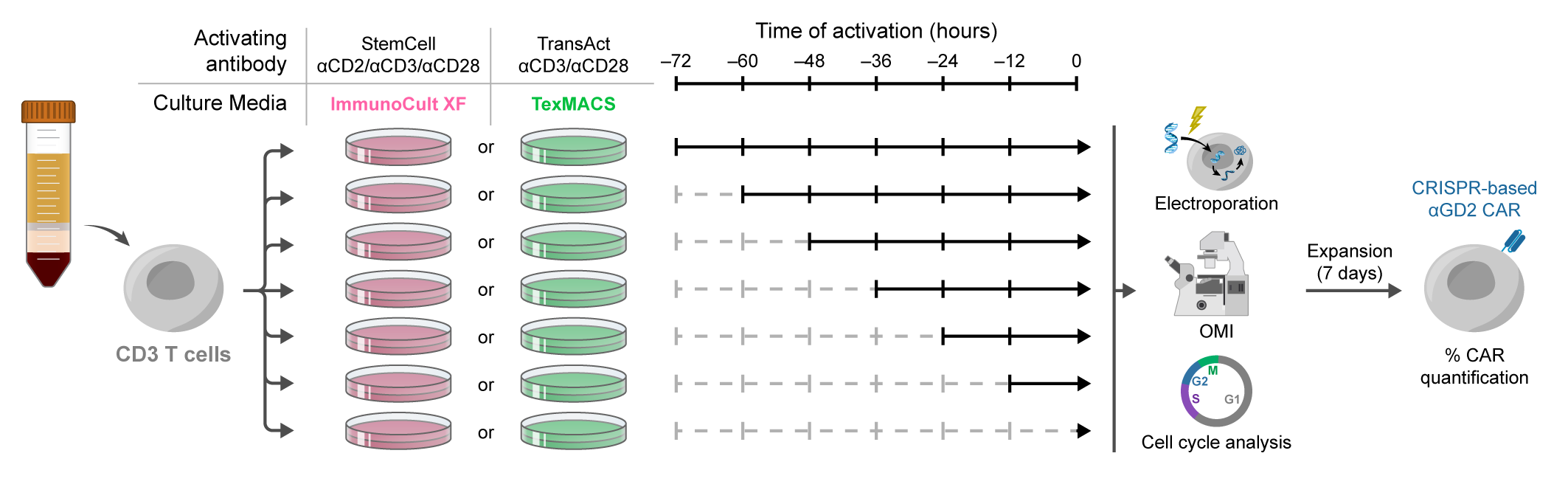

**Fig. S1. Activation time-course and conditions for anti-GD2 CAR T cell generation with CRISPR/Cas9.** CD3 T cells were isolated from 3 healthy donors and activated with StemCell αCD2/αCD3/αCD28 in ImmunoCult XF media (Imm) or TransAct αCD3/αCD28 in TexMACs media (Tex). Cells were divided into 7 groups with duration of activation ranging from 0 hours (quiescent T cells) to 72 hours prior to EP to introduce the anti-GD2 CAR transgene. When activated T cells were collected for EP, their metabolic features were characterized using OMI. Meanwhile, their cell cycle stage and proliferation capacity were assessed using flow cytometry of Hoechst 33342 and Ki-67 staining. These activated T cells were then electroporated to incorporate the anti-GD2 CAR transgene and expanded in corresponding culture media. Genome editing efficiency was determined after 7 days of expansion with GD2 CAR antibody using flow cytometry.

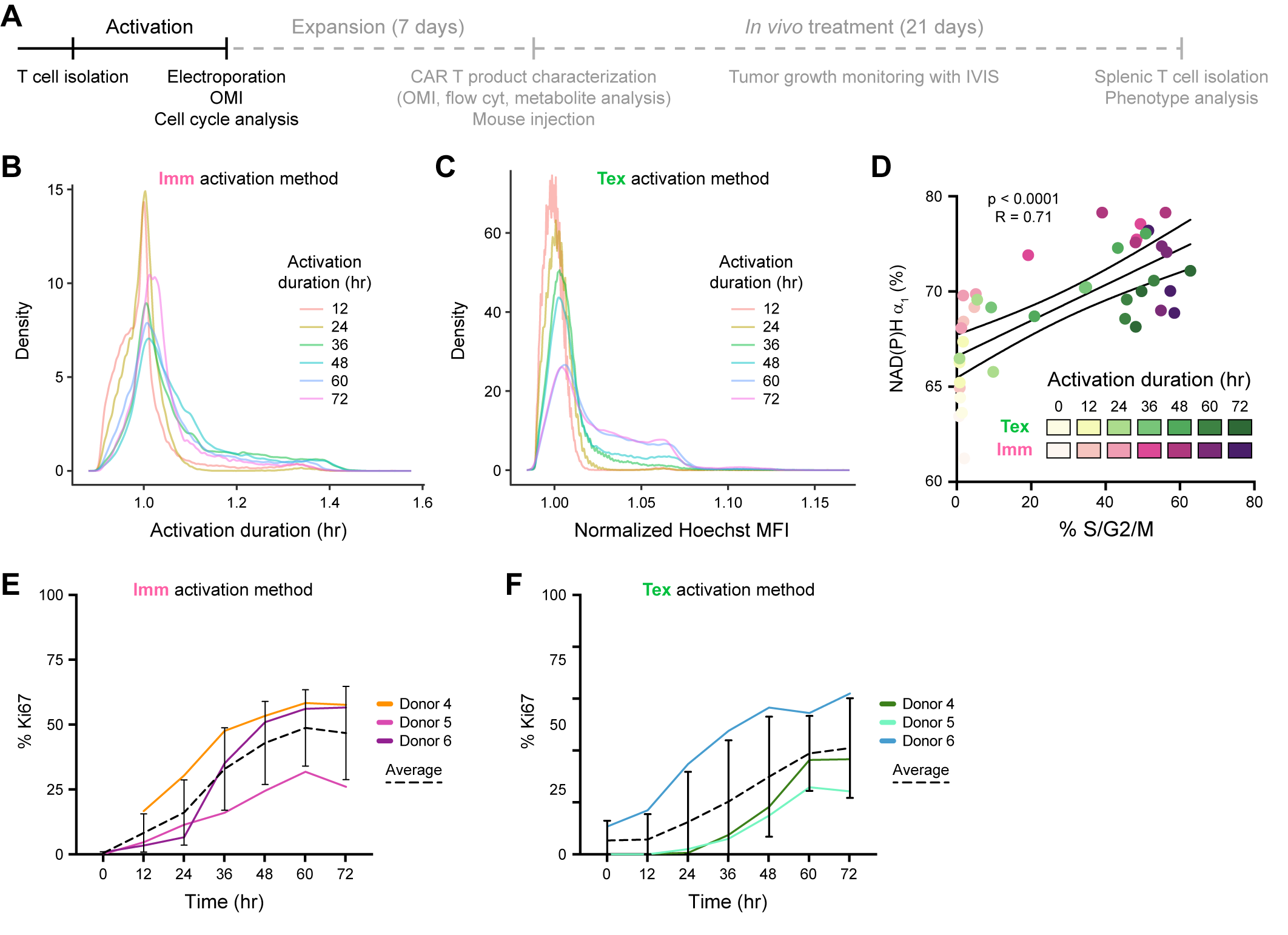

**Fig. S2. T cells progressed through cell cycle and proliferated upon activation. (A)** Experimental timeline**.** **(B-C)** T cells progressed through cell cycle upon activation. Density plots of Hoechst MFI in T cells activated with **(B)** Imm and **(C)** Tex methods. For each donor, Hoechst MFI was normalized to the average of 12-hour-activated group. n = 393,999 cells and 370,802 cells for Imm and Tex activation methods, respectively. **(D)** Correlation between NAD(P)H α_1_ and cell cycle stage (% cells in S/G2/M phase) at EP. Each dot represents one sample average, color coded based on the method of activation (Imm or Tex) and duration of activation (0–72 hours). n= 42 samples, Pearson R analysis. **(E-F)** T cell proliferation (% Ki-67^+^) following activation with **(E)** Imm method or **(F)** Tex method. n = 3 donors, Brown-Forsythe and Welch ANOVA test with Dunnett’s post hoc test for multiple comparisons. Bars are mean ± SD. Color lines connected donor-match averages.

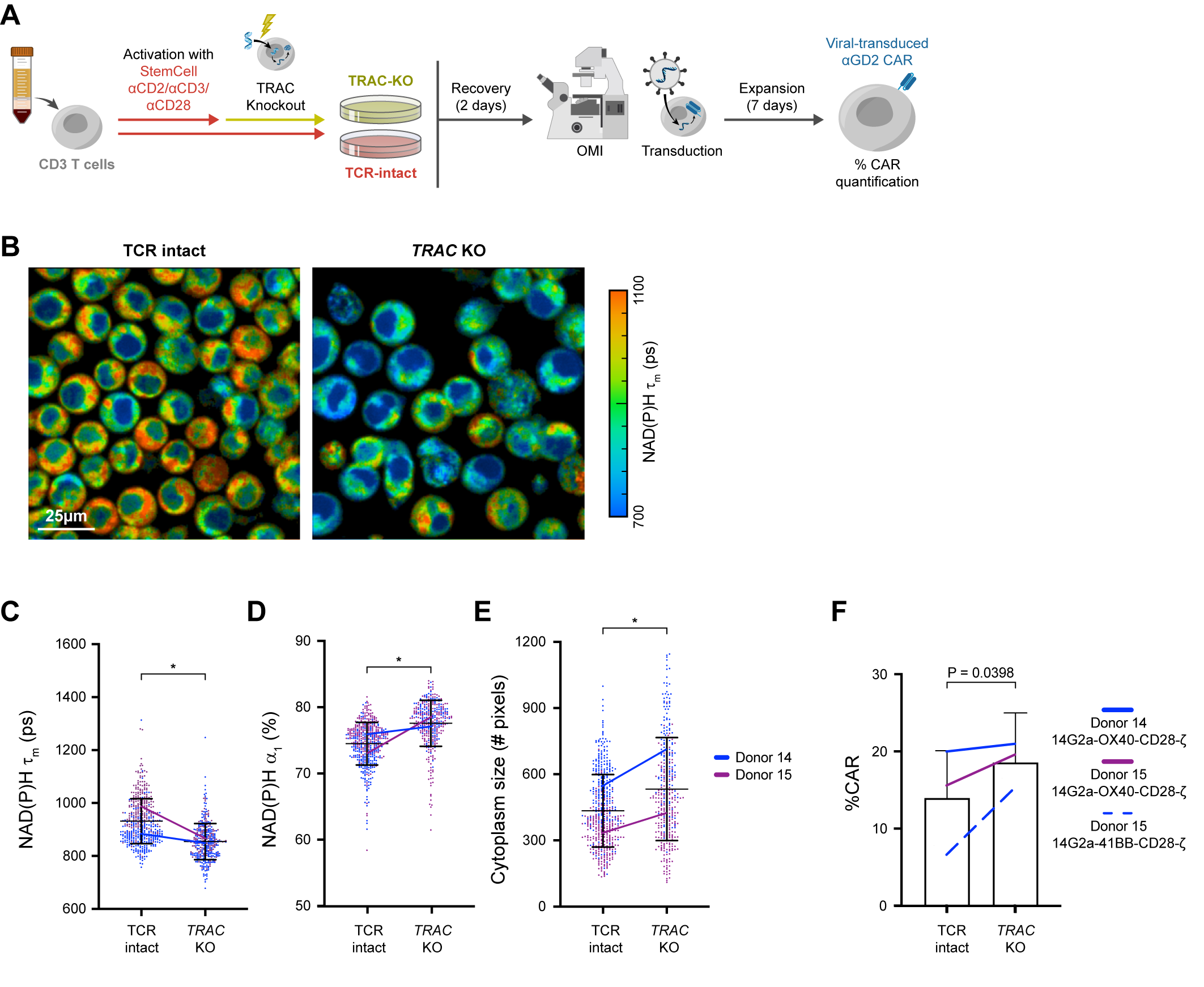

**Fig. S3. T cell metabolism at viral transduction timepoint correlated with transduction efficiency.** **(A)** Experimental setup. T cells were isolated from peripheral blood of two healthy donors and activated with Imm method. Activated CD3 T cells underwent electroporation with Cas9 ribonucleoproteins targeting the human *TRAC* locus to knockout the T cell receptor (*TRAC* KO) 2 days prior to transduction with retrovirus to express anti-GD2 CAR receptor using two constructs (OX40-CD28-CAR or 41BB-CAR) as previously described^9^. OMI was performed immediately before transduction. Transduction efficiency was quantified as %CAR^+^ after 7 days of expansion. **(B)** Representative NAD(P)H τ_m_ images of TCR-intact and TRAC KO T cells at transduction. OMI parameters including **(C)** NAD(P)H τ_m_, **(D)** NAD(P)H α_1_, and **(E)** cytoplasm size of cells with intact T cell receptor (TCR-intact) and *TRAC* KO T cells at transduction. n = 374-480 cells/group from 2 independent donors, Mann-Whitney test. **(F)** Transduction efficiency (% CAR) of TCR-intact and TRAC KO cells at the end of the manufacturing process. n = 17 samples from 2 donors and 2 anti-GD2 CAR constructs, unpaired T test. Bars are mean ± SD. Color lines connected donor-match averages. * p < 0.0001.

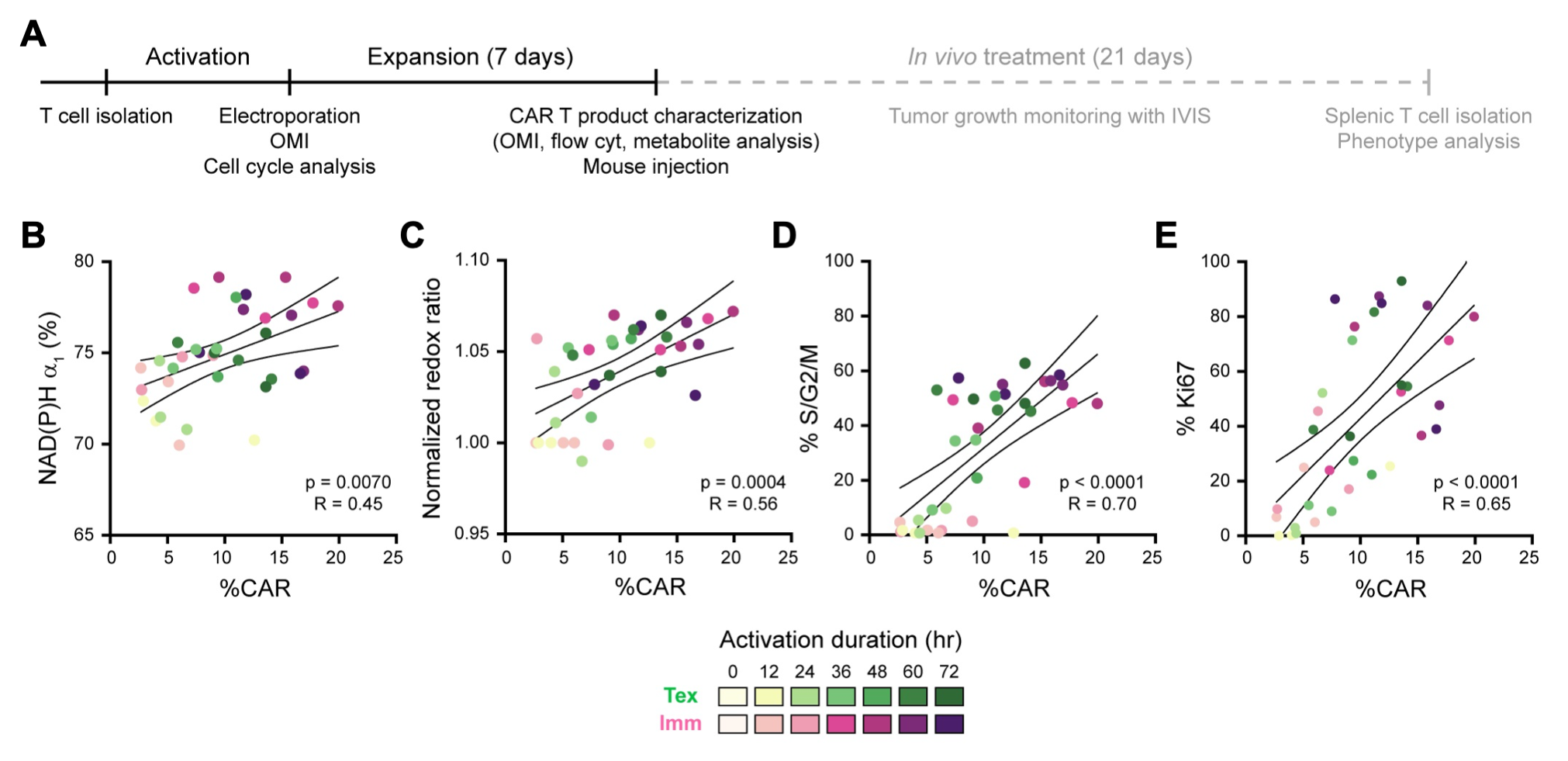

**Fig. S4.** **Cell characteristics at EP correlated with genome editing efficiency.** **(A)** Experimental timeline. **(B-E)** Correlation between cell characteristics (**(B)** NAD(P)H α_1_, **(C)** normalized redox ratio, **(D)** %S/G2/M and **(E)** %Ki-67^+^) at EP and genome editing outcome (% CAR). Each dot represents one sample average, color coded based on the method of activation (Imm or Tex) and duration of activation (0–72 hours). n= 36 samples, Pearson R analysis. Quiescent T cells with 0-hour activation duration did not undergo genome editing.

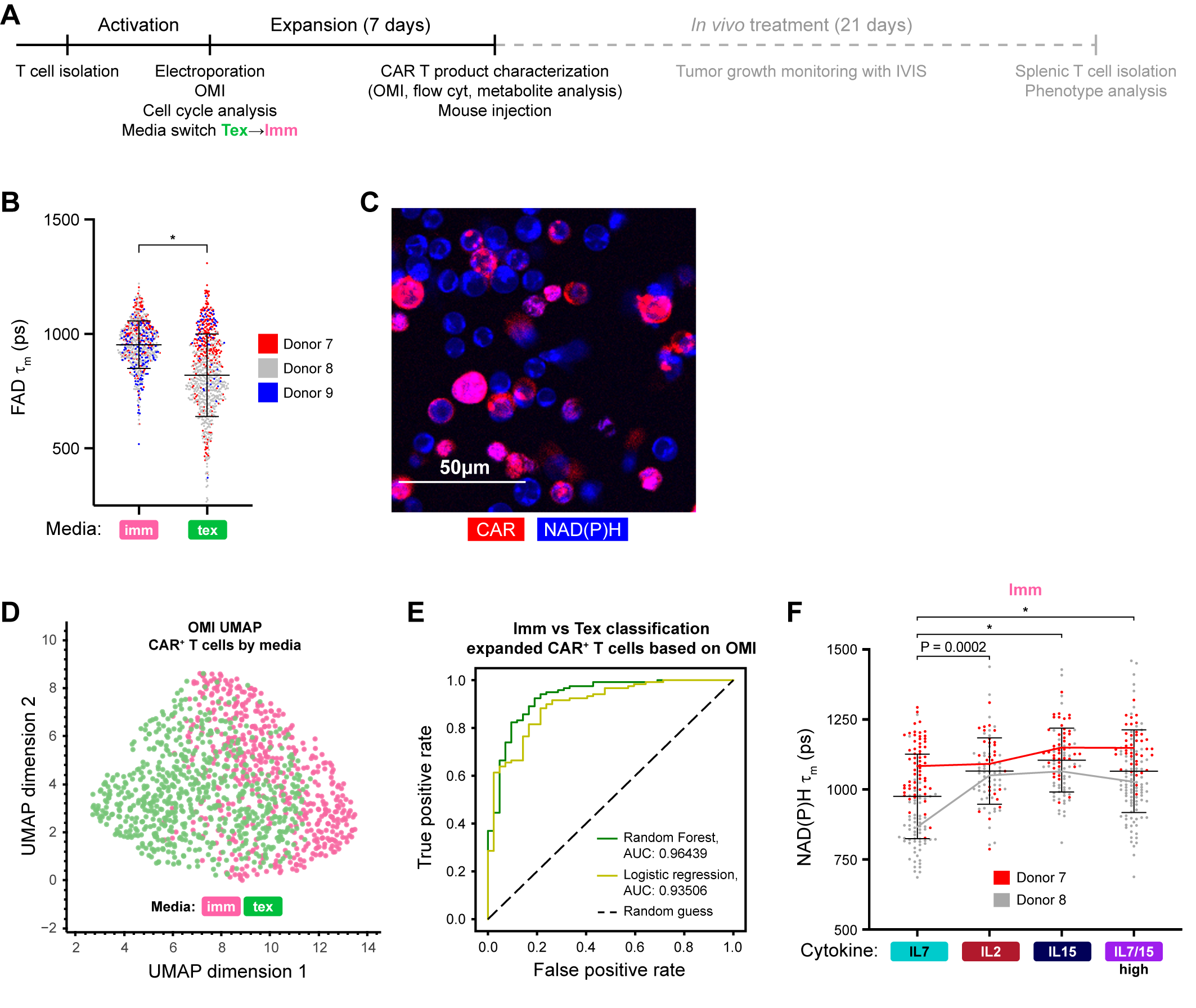

**Fig. S5. OMI revealed metabolic differences in CAR T cells expanded in Imm and Tex media (A)** Experimental timeline. **(B)** Quantification of FAD mean lifetime (FAD τ_m_) of CAR T cells expanded in either ImmunoCult XF or TexMacs media. n = 666-751 cells/condition across 3 independent donors, Mann-Whitney test. **(D-F)** OMI metabolic profiles of CAR^+^ T cells differed based on expansion media. **(D)** Representative image of CAR+ T cells identified based on PerCP conjugated GD-2 CAR antibody (red). **(E)** UMAP based on 13 OMI parameters of CAR+ T cells expanded in Imm or Tex media. **(F)** ROC curves and AUCs of three models based on OMI parameters (Table 1) to classify CAR+ T cells by expansion media (Imm vs. Tex); n = 494 cells for training (80%), n = 123 cells for testing (20%). **(G)** Quantification of NAD(P)H τ_m_ from CAR T cells expanded in ImmunoCult XF media supplemented with 500U/mL IL-2 compared to other cytokine combinations, n=83-129 cells/condition from 2 donors, ANOVA with Kruskal-Wallis test. * p < 0.0001.

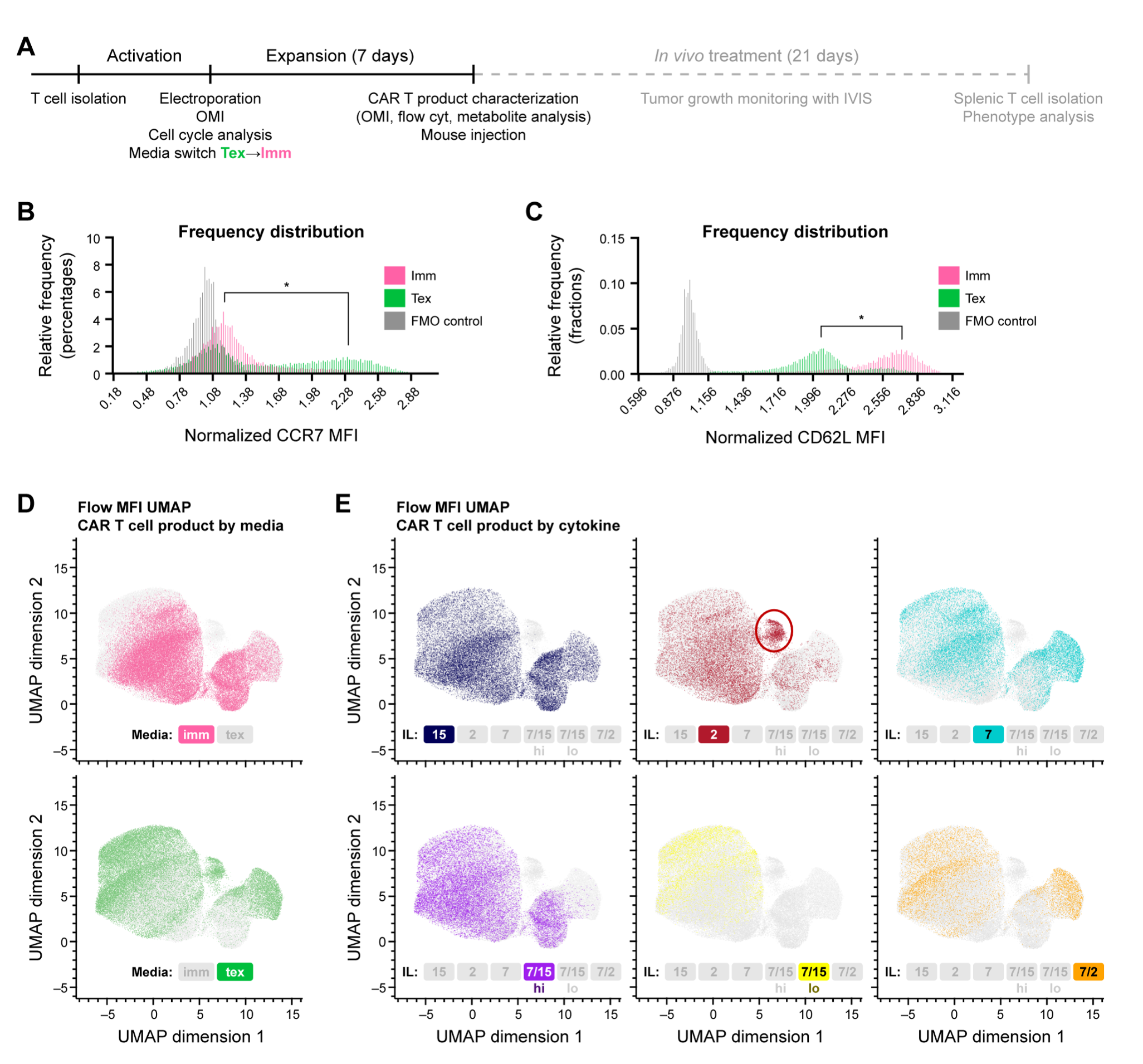

**Fig. S6. CAR+ T cells expanded in Imm and Tex media showed different phenotypes.** **(A)** Experimental timeline. MFIs of **(B)** CCR7 and **(C)** CD62L of CAR+ T cells expanded in either Imm or Tex media supplemented with different cytokine cocktails, normalized by fluorescence-minus-one (FMO) controls. (n = 87317 cells from 3 independent donors). UMAP based on Euclidean distances of MFIs of 6 surface markers (CD27, CD45RO, CD62L, CD28, CD45RA and CCR7) of CAR^+^ T cells expanded in Imm and Tex media, color coded based on **(D)** culture media or **(E)** supplemented cytokines. Dots are CAR^+^ T cells. n= 87317 cells from 3 independent donors. * p < 0.0001.

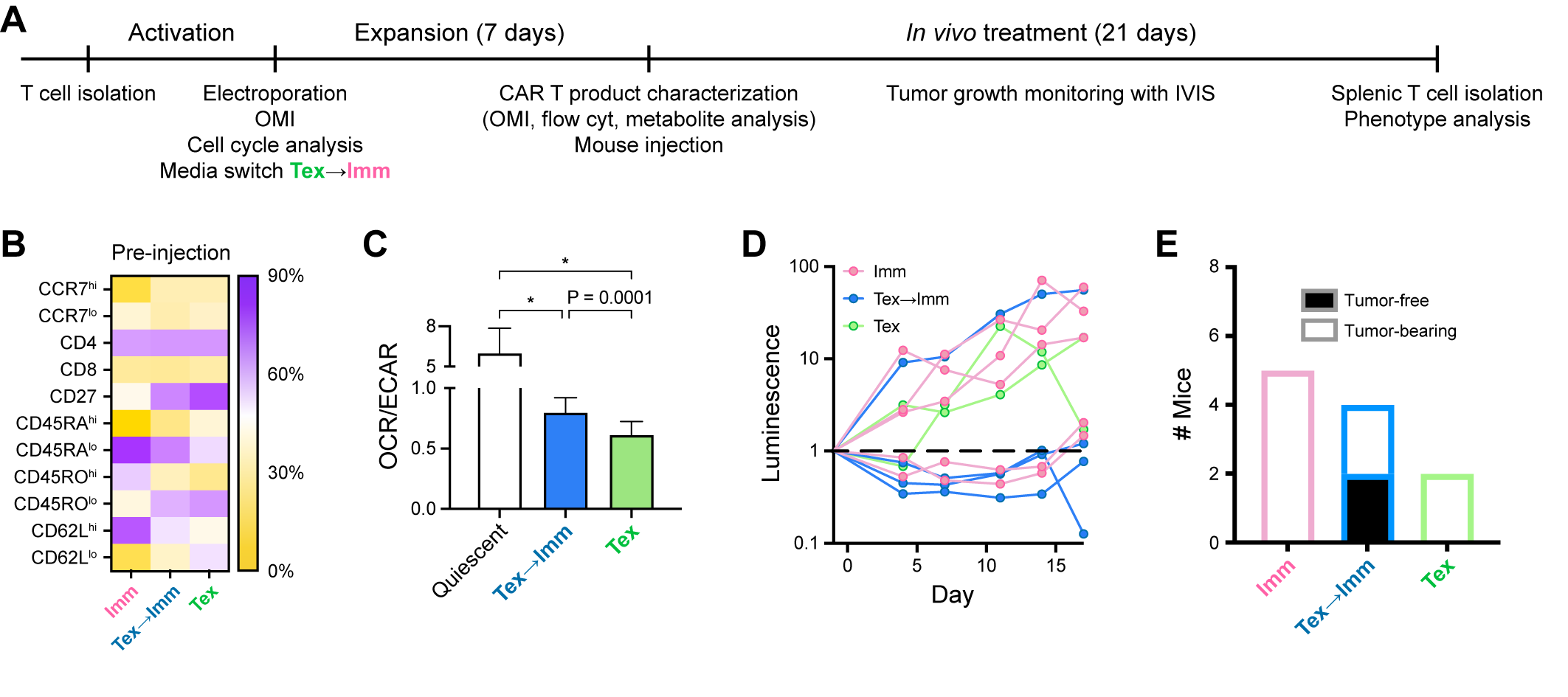

**Fig. S7. CAR T cell phenotypes and metabolic profile before *in vivo* treatment course and *in vivo* treatment response. (A)** Experimental timeline. **(B)** Phenotypes of CAR+ T cells expanded in Imm, Tex🡪Imm, and Tex conditions based on expression of 7 surface markers. **(C)** Extracellular flux analysis of baseline oxygen consumption rate (OCR) and extracellular acidification rate (ECAR) from CD3 T cells that were unactivated (quiescent) or activated and expanded in Imm or Tex🡪Imm condition. n=9-18 samples/condition Brown-Forsythe and Welch ANOVA test with Dunnette T3 post hoc test for multiple comparisons against Tex🡪Imm group. **(D)** Fold change in tumor flux (measured with IVIS imaging) in NSG tumor-bearing mice after CAR T cell treatment. Dash line represents no change in tumor flux (fold change = 1). **(E)** CAR T treatment outcomes on day 17 post treatment. * p < 0.0001.

**Supplementary Tables**

| Variable | ΔNAD(P)H τ_m_ | | ΔNAD(P)H α_1_ | |
| --- | --- | --- | --- | --- |
| Activation method | Imm | Tex | Imm | Tex |
| Donor 1 | 2.61 | 1.56 | 3.76 | 2.50 |
| Donor 2 | 3.55 | 1.66 | 5.53 | 3.35 |
| Donor 3 | 2.18 | 1.57 | 4.64 | 2.47 |

**Table S1. Glass’s deltas (**Δ) **showing maximal effect sizes of Imm and Tex activation methods on T cell metabolism (NAD(P)H τ_m_ and NAD(P)H α_1_).** For each donor and activation method, Glass’s deltas were calculated based on NAD(P)H τ_m_ and NAD(P)H α_1_ at the time point (12, 24, 36, 48, 60, or 72 hours of activation duration) of maximal differences compared to donor-matched quiescent T cells (0hr activation duration) in respective media (Imm or Tex).
